# Comprehensive mapping of the human cytokine gene regulatory network

**DOI:** 10.1101/2020.05.05.079657

**Authors:** CS Santoso, Z Li, S Lal, S Yuan, KA Gan, LM Agosto, X Liu, S Carrasco Pro, JA Sewell, A Henderson, MK Atianand, JI Fuxman Bass

## Abstract

Proper cytokine gene expression is essential in development, homeostasis, and immune responses. Studies on the transcriptional control of cytokine genes have mostly focused on highly researched transcription factors (TFs) and cytokines, resulting in an incomplete portrait of cytokine gene regulation. Here, we use enhanced yeast one-hybrid (eY1H) assays to derive a comprehensive network comprising 1,380 interactions between 265 TFs and 108 cytokine gene promoters, greatly expanding the known repertoire of TF-cytokine gene interactions. We found an enrichment of nuclear receptors and confirmed their role in cytokine regulation in primary macrophages. Additionally, we used the eY1H-derived network as a framework to identify pairs of TFs that synergistically modulate cytokine gene expression, and to identify novel TF-cytokine regulatory axes in immune diseases and immune cell lineage development. Overall, the eY1H data provides a rich resource to study cytokine regulation in a variety of physiological and disease contexts.

## Introduction

Transcriptional regulation of cytokine genes plays a central role in the development of the immune system, responses to pathogens, and inflammation (Griffith et al., 2014; Medzhitov and Horng, 2009). Indeed, dysregulation of cytokine expression, caused by mutations in cytokine gene regulatory regions, transcription factors (TFs), and genes belonging to upstream signaling pathways that impinge on cytokine transcriptional control regions, have been associated with autoimmune diseases, immunodeficiency, and susceptibility to infections (Turner et al., 2014). Thus, identifying the repertoire of TFs that regulate each cytokine gene is central to understanding the mechanisms that control immune responses, which will aid in the design of novel therapeutic strategies to modulate cytokine expression in immune diseases.

Research conducted for more than three decades has identified 170 TFs that bind to the transcriptional control regions and/or regulate 95 (of ∼140) human cytokine genes (Carrasco Pro et al., 2018). This includes TFs that are activated by pathogen signals (*e.g.*, NF-κB, AP-1, and IRFs), stress signals (*e.g.*, HIF1A, TP53, and HSF1), cytokine signals (*e.g.*, STATs and NF-κB), as well as lineage factors (*e.g.*, SPI1 and CEBPA). However, analysis of this literature-derived human cytokine protein-DNA interaction (PDI) network suggests that this network is largely incomplete. For instance, no PDIs have been reported for nearly 30% of the cytokine genes, and new TFs and PDIs in the cytokine PDI network are still being reported at a rate of 6.6 TFs and 35 PDIs per year (Carrasco Pro et al., 2018). More importantly, most PDIs reported in the literature correspond to highly studied TFs and cytokines. Whether this bias has a biological basis or is due to research trends that arise from reagent availability or “fashionable” TFs and cytokines, remains to be determined. Taken together, these facts suggest that comprehensive and unbiased screens are needed to delineate a more complete cytokine PDI network.

Different approaches have been used to identify PDIs in a high-throughput manner. Chromatin immunoprecipitation followed by next generation sequencing (ChIP-seq) has been widely used to identify the DNA regions where a TF binds. Although ChIP-seq has provided a wealth of information regarding TF genomic occupancy, ChIP-seq data are still largely incomplete. Only 30% of human TFs have been tested due to the lack of ChIP-grade anti-TF antibodies, and ChIP-seq has only been performed in a limited number of cell types and conditions, mostly in non-immune cell lines in basal conditions (Encode Project Consortium, 2012; Yevshin et al., 2017). Another approach to infer PDI networks involves integrating ATAC-seq or DNase-seq footprinting data with DNA-binding site preferences and expression levels of TFs (Pokrovskii et al., 2019). However, these analyses are challenging for multiple reasons: 1) transposition and cleavage preferences have to be distinguished from true DNA protection due to TF binding, 2) DNA binding specificities have not been determined for approximately half of human TFs, 3) similar DNA binding specificities between TF paralogs often lead to multiple TFs being predicted to bind a protected site, and 4) multiple cells need to be assayed in different conditions due to differences in chromatin landscape between cells.

Enhanced yeast one-hybrid (eY1H) assays is a powerful PDI mapping method that tests the binding of hundreds of TFs to a DNA region of interest (*e.g.*, a cytokine promoter) in the milieu of the yeast nucleus (Reece-Hoyes et al., 2011; Shrestha et al., 2019a). This approach circumvents the requirement of antibodies in ChIP-seq and testing in multiple cells and conditions in ChIP-seq, ATAC-seq, and DNase-seq. eY1H assays constitute a robust system to identify PDIs given that it uses a dual reporter system, each interaction is tested in quadruplicate, and positive interactions are identified by integrating image analysis software and manual curation (Fuxman Bass et al., 2015; Reece-Hoyes et al., 2013). More importantly, PDIs identified by eY1H assays display a 30-70% validation rate in human cells and living animals, which is similar to the validation rate of ChIP-seq interactions (Fuxman Bass et al., 2016; Fuxman Bass et al., 2015; Whitfield et al., 2012). We previously developed a high-throughput eY1H pipeline that interrogates the binding of approximately two-thirds (1,086 of ∼1,600) of all human TFs to defined regulatory regions, which we used to identify TFs that bind to human developmental enhancers, noncoding genetic variants, and repetitive DNA elements (Fuxman Bass et al., 2015; Shrestha et al., 2019b).

Here, we used eY1H assays to interrogate the binding of 1,086 human TFs to human cytokine promoters, from which we derived a comprehensive cytokine PDI network comprising 1,380 PDIs between 265 TFs and 108 cytokine promoters, substantially expanding the current literature-derived cytokine PDI network. Using orthogonal assays, we observed a validation rate similar to PDIs reported in the literature or identified by ChIP-seq, and we experimentally confirmed a regulatory function for 175 eY1H-derived PDIs in mammalian cell lines or primary macrophages. Finally, we leveraged the eY1H network as a framework to identify (1) pairs of TFs that can be targeted with commercially-available drugs to synergistically modulate cytokine production, and (2) novel TFs and TF-cytokine regulatory axes in the pathogenesis of immune diseases and in the development of immune cell lineages. Altogether, these studies demonstrate the use of the eY1H-derived network as a powerful resource to study cytokine regulation in a variety of cellular and disease contexts.

## Methods

### Generation of CytReg v2

To expand the list of literature-reported physical and regulatory PDIs between TFs and cytokine genes annotated in CytReg (https://cytreg.bu.edu)(Carrasco Pro et al., 2018), in February 2019 we searched for papers mentioning TF-cytokine pairs and key words “bind”, “direct”, “target”, “control”, “regulate”, “activate”, “repress”, or “suppress”. The resulting papers were manually curated to determine whether direct experimental evidence for the PDI was provided. Additionally, the species, type of assay, and functional activity (activating or repressing) when provided, were also annotated. In total, 160 new literature-derived interactions, and 1,380 eY1H-derived interactions were added to generate CytReg v2 (https://cytreg.bu.edu/search_v2.html) (**Tables S1** and **S3**).

### Bias in literature-reported PDIs

The number of publications in PubMed for TFs and cytokines were retrieved from NCBI’s Gene database on August 16, 2019. TFs and cytokines were ranked by the number of publications and then partitioned into equal-sized bins. The number of interactions in the literature-derived or the eY1H-derived PDI network between TFs and cytokines in each pair of bins in the two-dimensional matrix were counted. The transcript per million (TPM) expression levels of TFs in 20 immune cell-types was obtained from data published by the Blueprint Epigenome Consortium (Stunnenberg et al., 2016)(http://dcc.blueprint-epigenome.eu). The maximum expression of each TF across the 20 immune cell-types was used to determine the average expression of TFs in immune cells in each bin. The association between TFs and immune diseases was obtained from genome-wide association studies (GWAS) downloaded from the NHGRI-EBI Catalog (MacArthur et al., 2017) on July 27, 2017 (**Table S2**). Terms describing autoimmune diseases and susceptibility to infections were used to determine whether a TF is associated with an immune disease. The association between TFs and immune phenotypes was determined from knockout mouse studies reported in the Mouse Genome Informatics (MGI) database (Eppig et al., 2017) on March 3, 2019 (**Table S2**). All terms classified as an “immune system phenotype” in MGI were used to determine whether a TF is associated with an immune phenotype.

### Enhanced yeast one-hybrid assays

Enhanced yeast one-hybrid (eY1H) assays were performed as previously described (Reece-Hoyes et al., 2011; Shrestha et al., 2019a) to detect PDIs between TFs and cytokine gene promoters by mating ‘TF-prey strains’ with ‘DNA-bait strains’. DNA-bait strains for 112 human cytokine promoters (∼2kb upstream of the transcription start sites) were obtained by PCR amplification (**Table S12**) using Platinum *Taq* DNA Polymerase High Fidelity (ThermoFisher) from a pool of human genomic DNA (Clonetech). PCR products were Gateway cloned into the pDONR P1-P4 vector and entry clones were confirmed by Sanger sequencing. Then DNA-baits were Gateway cloned upstream of two reporter genes (*HIS3* and *LacZ*) and both reporter constructs were integrated into the yeast genome to generate chromatinized DNA-bait strains (Deplancke et al., 2006a; Deplancke et al., 2006b). Yeast DNA-bait strains were confirmed by Sanger sequencing. DNA-bait strains were mated with an array of 1,086 TF-prey yeast strains expressing human TFs fused to the yeast Gal4 activation domain (AD) (Fuxman Bass et al., 2015). Matings were performed using a Singer Rotor robotic platform that manipulates yeast strains in a 1,536-colony format. An interaction was detected when a TF-prey binds the DNA-bait and the AD moiety activates reporter gene expression, allowing the mated yeast to grow on media lacking histidine, overcome the addition of 3-amino-triazole (3AT), a competitive inhibitor of the His3p enzyme, and convert the colorless X-gal into a blue compound. Each interaction was tested in quadruplicate and interactions were considered positive if at least two of the four mated colonies tested positively (>90% of detected interactions tested positively for all four colonies).

Images of mated colonies on plates lacking histidine and containing 3AT and X-gal were processed using the Mybrid web-tool to detect positive interactions (Reece-Hoyes et al., 2013). Positive interactions detected by Mybrid were then manually checked and curated. False positives detected by Mybrid, which typically occur on plates with uneven background, were removed. False negatives missed by Mybrid, which typically occur when baits exhibit high background reporter gene expression or when interactions occur next to very strong positives, were included. A total of high-quality 1,380 PDIs between 265 TFs and 108 cytokine promoters were included in the final dataset (**Table S3**). The eY1H interactions from this publication have been submitted to the IMEx (http://www.imexconsortium.org) consortium through IntAct [X] and assigned the identifier IM-27908 (Orchard et al., 2014).

### Overlap between ChIP-seq and eY1H interactions

ChIP-seq peak coordinates were downloaded from the ENCODE Project (Encode Project Consortium, 2012) and GTRD database (Yevshin et al., 2017) on August 27, 2018. Comparison between eY1H and ChIP-seq interactions were limited to 130 TFs that were both detected by eY1H and tested by ChIP-seq. ChIP-seq peaks were assigned to a particular promoter if the midpoint of the peak was located within the promoter sequence. Multiple ChIP-seq peaks mapping to the same promoter were counted as one TF-promoter interaction. We compared the PDIs derived from ChIP-seq to the PDIs observed in the eY1H network or in 10,000 randomized versions of the eY1H data to determine the significance of the overlap.

### Overlap between TF binding sites and eY1H interactions

Position weight matrices (PWMs) for 194 human TFs in the eY1H network were downloaded from CIS-BP (Weirauch et al., 2014). The presence of TF binding sites within the cytokine promoter baits were determined based on PWM matches. Each position within the cytokine promoters was scored by calculating the sum of logs for each PWM and significant scores were determined using TFM P-value (Touzet and Varre, 2007). Scores above the TFM score threshold for P-value < 10^−4^ for a given PWM were considered significant hits and thereby predicted PDIs. To determine statistical significance of the overlap between eY1H and predicted PDIs, we compared the predicted PDIs to 10,000 randomized versions of the eY1H network.

### Network randomizations

To determine the significance of overlap between the eY1H network and ChIP-seq and TF binding site datasets, the eY1H network was randomized by edge switching while preserving both overall network topology and individual node degree. Briefly, edge pairs (e.g., A-B and C-D) chosen at random from the network were swapped (i.e., A-D and C-B)(Martinez et al., 2008). Edges were switched only if neither of the new edges were already present in the network. Each random network was generated from >20,000 edge switching events. We generated 10,000 random networks to determine PDI overlap with the ChIP-seq and TF binding site datasets. Based on the overlap with each of the 10,000 random networks, a Z-score was determined to calculate the P-value of the overlap corresponding to the eY1H-derived network.

### Luciferase assays

Cytokine-promoter baits were cloned upstream of the firefly luciferase reporter gene in a Gateway compatible pGL4.23[luc2/minP] vector (Fuxman Bass et al., 2015). TFs were cloned into the Gateway compatible pEZY3 vector (Addgene) or pEZY3-VP160 vector such that TFs are fused to 10 copies of the VP16 activation domain (Carrasco Pro et al., 2018). To perform luciferase assays, HEK 293T cells were cultured in DMEM supplemented with 10% FBS and 1% Antibiotic-Antimycotic (100X) and plated in 96-well white opaque plates (∼1 x 10^4^ cells/well). 24 hours later, cells were transfected using Lipofectamine 3000 (Invitrogen) according to the manufacturer’s protocol, with 80 ng of a TF-pEZY3 or TF-pEZY3-VP160 plasmid, 20 ng of a cytokine promoter-pGL4.23[luc2/minP] firefly luciferase plasmid, and 10 ng of the renilla luciferase plasmid as a transfection normalization control. An empty pEZY3 or pEZY3-VP160 plasmid co-transfected with the corresponding recombinant firefly luciferase plasmid were used as negative controls. 48 hours after transfection, firefly and renilla luciferase activities were measured using the Dual-Glo Luciferase Assay System (Promega) according to the manufacturer’s protocol. Non-transfected cells were used to subtract background firefly and renilla luciferase activities, and then firefly luciferase activity was normalized to renilla luciferase activity in each well.

### TF knockdown in human PBMC-derived macrophages

Peripheral blood mononuclear cells (PBMCs) were isolated from de-identified human leukapheresis-processed blood (New York Biologics, Inc) by centrifugation through Lymphoprep (Stem Cell Technologies.) density gradient. Purified PBMCs were resuspended in serum-free RPMI medium and plated in 12-well or 6-well plates at a density of 5 × 10^6^ cells/ml. Cells were incubated at 37°C for 1–2 hours to allow binding of monocytes to the plates, then the medium and unbound cells were discarded and replaced with RPMI supplemented with 10% FBS, 10% human AB serum (Corning), 2 mM L-glutamine (Invitrogen), and 100U/ml of penicillin/streptomycin (Invitrogen). Monocytes differentiated into macrophages over 7 days at 37°C, with fresh medium replenished every 2–3 days. On day 8, cells were transfected using Lipofectamine 3000 according to the manufacturer’s protocol, with 50 pmol/ml of ON-TARGETplus SMART-pool siRNA (GE Dharmacon) to knockdown the respective TFs. On day 10, cells were treated with 10 ng/ml LPS for 4 hours and then harvested in TRIzol and analyzed by RT-qPCR. Each experimental condition was performed in three biological replicates.

### Generation of iBMDM stable cell lines and stimulation with ligands

HEK293T and immortalized bone-marrow-derived macrophages (iBMDMs) were cultured in DMEM supplemented with 10% FBS and 1% Pen-Strep under standard cell culture conditions. Control and gene-specific shRNAs (**Table S13**) were cloned in pLKO.1 lentiviral vector and sequences were verified by Sanger sequencing (Genewiz). Stable iBMDMs cell lines were generated as described previously (Atianand et al., 2016). In brief, HEK293T cells were co-transfected with pLKO.1 shRNA plasmid (500 ng) together with the packaging plasmids, psPAX (375 ng) and pMD2 (125 ng), using GeneJuice (Novagen). Culture supernatants containing viral particles were collected at 48 and 72 hours post transfection, pooled together and filtered through 0.45 mm PVDF membrane filter (Sigma). iBMDMs were transduced with viral particles in the presence of polybrene (8 mg/ml), and cells were selected with puromycin (3 ug/ml). iBMDMs were either stimulated with *E. coli* LPS (1 ug/ml) or infected with Sendai virus (200 HAU/ml) or *E. coli* strain DH5α (1 CFU/cell) for 6 hrs. RNA was extracted using Trizol and analyzed by RT-qPCR.

### RT-qPCR measurments

To measure the expression of TFs and cytokines in human PBMC-derived macrophages or mouse iBMDMs, total RNA was extracted using TRIzol (Invitrogen) and then purified using Direct-zol RNA MiniPrep kit (Zymo Research) including the DNAse I treatment step to remove contaminating DNA. cDNA was reverse-transcribed from RNA using random hexamers and M-MuLV reverse transcriptase (NEB). Primer sequences for qPCR were designed using Primer3 such that primers are located in different exons or in exon-exon junctions and checked for any off-targets using the NCBI Primer-BLAST tool. qPCRs were performed in two technical replicates using the PowerUp SYBR Green Master Mix (ThermoFisher Scientific) and primers listed (**Table S14**). Relative transcript abundances were calculated using the ΔΔC_t_ method and normalized to *RPL13A* and *GAPDH* mRNA levels for human and mouse genes, respectively.

### Nuclear receptor functional interactions

The functional interactions between nuclear receptors (NRs) and cytokine genes were obtained from the literature-reported network and the Nuclear Receptor Signaling Atlas (NURSA) Transcriptomine resource (Becnel et al., 2017) on July 26, 2019 (**Table S4**). Functional interactions reported from genetic perturbation studies (e.g., NR knockout, knockdown, and overexpression) and ligand-based studies (e.g., NR activation or inhibition using agonists or antagonists) were considered. Ligands were assigned to the corresponding nuclear receptors based on assignments in NURSA, DrugBank, and Tocris (**Table S4**).

### Synergistic potentiation of IL10 production in M2-polarized THP-1 cells

THP-1 monocytes were cultured in RPMI supplemented with 10% FBS, 0.05 mM 2-mercaptoethanol, and 1% Antibiotic-Antimycotic (100X), and differentiated into macrophages with 100 nM phorbol 12-myristate 13-acetate (PMA) for 72 hours, and then recovered in fresh media for 48 hours. To perform the functional assays, THP-1-derived macrophages were incubated with the respective NR agonists/antagonists for 15 minutes, and then polarized to M2 macrophages with 25 ng/ml IL4 and 25 ng/ml IL13 for 48 hours to promote IL10 production. The supernatants were collected and the amount of IL10 in the supernatants were measured using the Human IL10 ELISA MAX (Biolegend) kit according to the manufacturer’s protocol. NR agonists/antagonists: NR1I2 agonist SR12813 (Tocris), NR1I2 antagonist SPA70 (Axon), RXR agonist Bexarotene (Tocris), RXR antagonist HX531 (Sigma Aldrich), and VDR agonist Ercalcitriol (Tocris).

### TF and cytokine associations with immune disorders and lineage differentiation abnormalities

The association between TFs or cytokines and abnormalities in immune cell differentiation or inflammatory diseases was obtained from DisGeNET (Pinero et al., 2017), GWAS (MacArthur et al., 2017), or MGI (Eppig et al., 2017). Gene-disease associations in DisGeNET were downloaded on September 20, 2019, associations in the NHGRI-EBI GWAS Catalog v1.0.2 were downloaded on September 4, 2019, and associations in MGI were downloaded on June 24, 2019. To determine whether a TF or cytokine is associated with one of the 12 immune diseases explored, terms describing allergy, asthma, eczema/psoriasis, fatty liver disease, general inflammation, inflammatory arthritis, inflammatory bowel disease, multiple sclerosis, primary biliary cholangitis, HIV, and susceptibility to infection, were considered (**Table S5**). To determine whether a TF or cytokine is associated with abnormalities in immune cell differentiation, terms describing abnormal differentiation, abnormal proliferation, and abnormal cell count, were considered (**Table S8**).

### Bulk RNA-seq data and analysis

The non-alcoholic fatty liver disease data was downloaded from the GEO repository GSE126848 (Suppli et al., 2019) as FASTQ files, and then were trimmed using Cutadapt (v2.2) for low quality bases and adapters. The trimmed FASTQ was aligned to the Ensembl Human Reference Genome (GrCh38.98) using STAR (v2.7.3a) (Dobin et al., 2013), and then quantified using featureCounts from subread (v1.5.0) (Liao et al., 2014). The Immgen data was downloaded from the GEO repositories GSE122597 (Yoshida et al., 2019), GSE109125 (Yoshida et al., 2019), and GSE122108 (Gal-Oz et al., 2019). The The International Human Epigenome Consortium (Stunnenberg et al., 2016) data were downloaded from http://www.cell.com/consortium/IHEC. The genes with lower read counts than 10 and standard deviations as 0 were excluded from the analyses.

The differential expression analyses for all dataset were conducted using DESeq2 (version 1.24.0) (Love et al., 2014), and then the log2 fold change shrinkage to remove noise was conducted using the APEGLM algorithm (Zhu et al., 2019). The Pearson correlation between TFs and cytokines were conducted using the log transformed TPM of genes and the Benjamini-Hochberg method was used to perform multiple hypothesis testing correction. The visualizations of data and results were conducted using the R package ggplot2 (v3.2.1) and PRISM (v8.4.0).

### Single cell RNA-seq data and analysis

The HIV data was downloaded from the GEO repository GSE117397 (Farhadian et al., 2018). The Crohn’s disease data was downloaded from the GEO repository GSE134809 (Martin et al., 2019). The hematopoietic progenitor cell data was downloaded from the GEO repository GSE117498 (Pellin et al., 2019). Each dataset was merged using Seurat (v3) (Butler et al., 2018) and cells with less than 200 genes and more than 20 percent mitochondrial contents were removed. The Louvain community network clustering algorithm was used to cluster cells and the MAST algorithm (Finak et al., 2015) was used to find the top markers for each cluster. Canonical markers for immune cell types were used to find different immune cell types. For the HIV dataset, we used the following markers described in the original article (Farhadian et al., 2018) to identify dendritic cells, B cells, and T cells, in the single cell dataset (**Figure S2**):

Dendritic cells: *ITGAM, ITGAX, ANPEP, CD33, CD80, CD7, CD83, CD86, CD27, CD28, CLEC4C, THBD, NRP1*

B cells: *MS4A1, CD79A, CD79B, CD19*

T cells: *CD2, CD3D, CD3G, CD4, CD5, CD7, CD8A, IL2RA, CD27, CD28, PTPRC*

For the Crohn’s disease dataset, we used the following markers described in the original article (Martin et al., 2019) to identify intestinal epithelial cells, dendritic cells, macrophages, B cells, and T cells (**Figure S3**):

Intestinal epithelial cells: *REG1B, REG1A, MUC13, ARL14, AGR2*

Dendritic cells: *ITGAM, ITGAX, ANPEP, CD33, CD80, CD7, CD83, CD86, CD27, CD28, CLEC4C, THBD, NRP1*

Macrophages: *CXCL2, MRC1, CD163, CD68, CSF1R, CD86, CD209, CD33, TLR2, TREM1, MSR1, CD63, LILRB1, IDO1, ENG, CD40, TGFB1, CXCL9, CXCL10, GPNMB, IL10, CD80, IRF4, STAT6*

B cells: *MS4A1, CD79A, CD79B, CD19, CD70*

T cells: *CD2, CD3D, CD4, CD5, CD7, CD8A, IL2RA, CD27, CD28*

Pearson correlation analyses were conducted for interactions in which the TF and the cytokine were detected in at least 25% of cells, and the Benjamin Hochberg (‘BH’) method was used to adjust the P-value from correlation analyses and identify significantly correlated (FDR < 0.25) TF-cytokine pairs (**Tables S6, S7**, and **S10**).

### Pseudo-time series analysis

The R package monocle2 (v2.4.0) (Trapnell et al., 2014) was used to conduct the pseudo-time series analysis. Briefly, a differential expression analysis was performed to identify the top significantly differentially expressed genes (FDR < 0.05) between healthy and disease conditions to build the disease trajectory, and then each single cell was assigned a numeric pseudotime value and then ordered along the disease trajectory (**Figures S2** and **S3**). TFs and cytokines are included in the analyses if their expressions were detected in at least 10% of cells within a cluster and are significantly differentially expressed (FDR < 0.25) along the pseudo-temporal relationship among cells.

## Results

### Biases and completeness of the literature-derived cytokine PDI network

We previously generated a literature-derived cytokine PDI network, in which biophysical (*i.e.*, determined from binding assays – ChIP or EMSA) and functional (*i.e.*, determined from reporter assays, or TF knockdown, knockout, or overexpression experiments) PDIs between TFs and cytokines genes were manually curated (Carrasco Pro et al., 2018). We have updated this network to include an additional 160 PDIs from the literature, expanding the network to 899 PDIs in human, 740 PDIs in mouse, and 73 PDIs in other species (**Table S1**). Although this literature-derived network constitutes the most comprehensive cytokine PDI network to date, we found marked research biases skewing the coverage of the network. When TFs and cytokines were ranked by the number of publications in PubMed and then partitioned into equal-sized bins, we found that nearly half of the reported PDIs were between the 10% most highly cited TFs and the 20% most highly cited cytokine genes (**Figure 1A**). Although an argument can be made that highly cited TFs and cytokines could have more pleiotropic roles, it is unlikely that lesser studied cytokines would be regulated by fewer or no TFs. Additionally, while the number of PDIs sharply decreases after the first TF bin, the expression levels of TFs in immune cells and the fraction of TFs associated with immune diseases in genome-wide association studies (GWAS) or immune phenotypes in knockout (KO) mouse studies (MGI) show a more gradual decrease across the TF bins (**Figure 1B** and **Table S2**). Taken together, these observations suggest that the skewed PDI distribution reflects a research bias towards highly studied TFs and cytokines.

**Figure 1.**
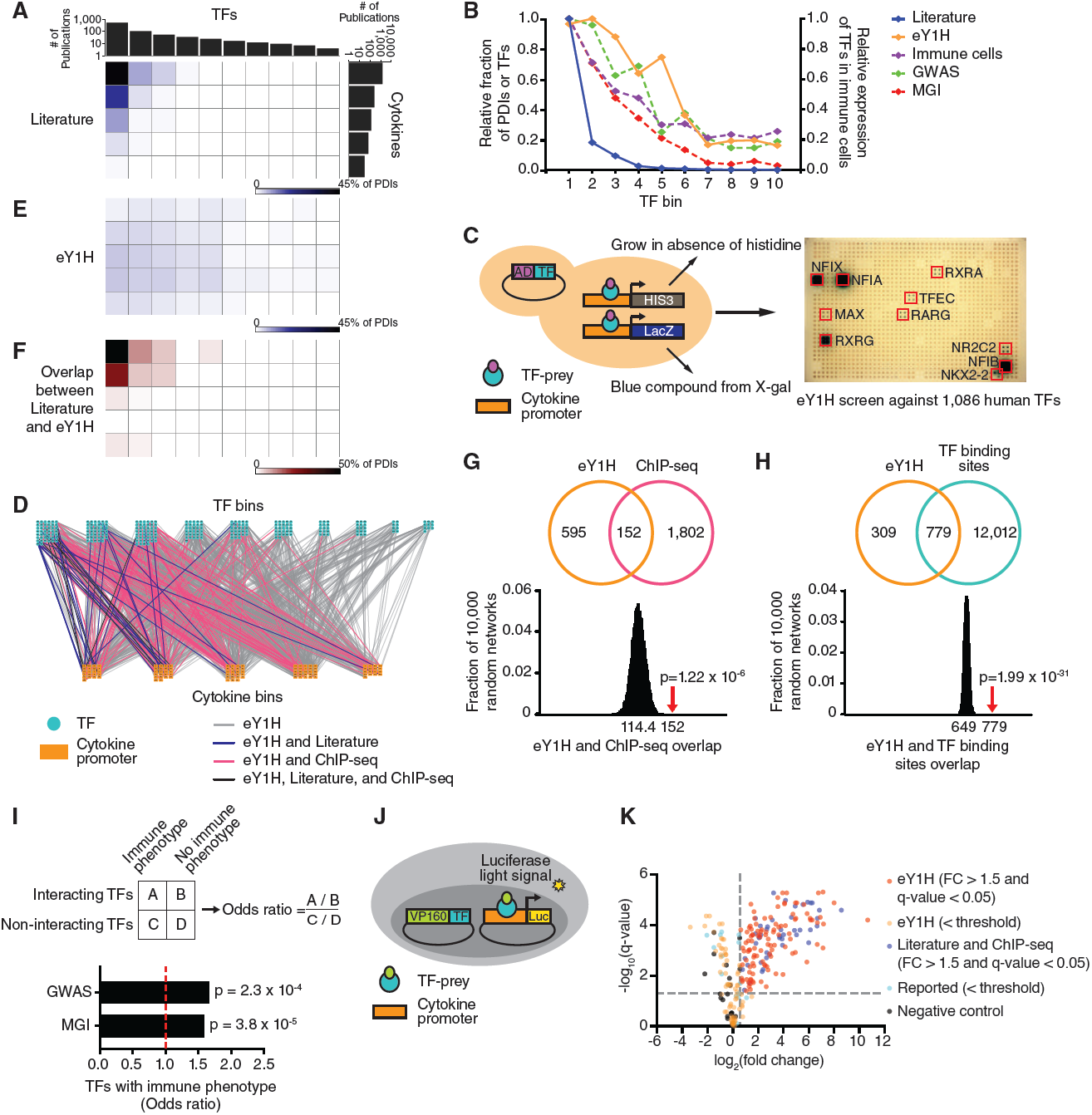
Cytokine PDI network. (A, E) Distribution of literature and eY1H PDIs based on the number of publications of TFs and cytokine genes. TFs and cytokine genes were ranked by the number of publications in PubMed and then partitioned into equal-sized bins. The intensity of blue in each matrix cell indicates the corresponding percentage of PDIs in the literature (A) or the eY1H network (E). (B) For each TF bin, the graph depicts the relative fraction of PDIs in the literature- or in the eY1H-derived network, the average expression of TFs in immune cells (Blueprint Epigenome), and the relative fraction of TFs associated with immune diseases in humans (GWAS) or immune phenotypes in KO mice (MGI). (C) Schematic of eY1H assays. A cytokine promoter (DNA-bait) is cloned upstream of two reporter genes (*HIS3* and *LacZ*) and integrated into the yeast genome. The resulting yeast DNA-bait strain is mated with a collection of yeast TF-prey strains harboring human TFs fused to the Gal4 activation domain (AD). Positive PDIs are determined by the ability of diploid yeast to grow in the absence of histidine, overcome the His3p competitive inhibitor 3AT, and turn blue in the presence of X-gal. Each PDI is tested in quadruplicate. Red boxes indicate positive interactions. (D) The eY1H cytokine PDI network comprises 1,380 PDIs (edges) between 265 TFs (turquoise nodes) and 108 cytokine gene promoters (orange nodes). TFs and cytokines are binned by the number of publications in PubMed. The eY1H PDIs reported in the literature (blue), ChIP-seq (pink), or both (purple) are indicated. (F) Overlap between literature and eY1H PDIs. The intensity of red indicates the percentage of eY1H PDIs reported in the literature. (G and H) eY1H PDIs significantly overlap with the occurrence of ChIP-seq peaks (G) and known TF binding sites (H) in the corresponding cytokine promoters. The Venn diagrams indicate the number of overlapping interactions. The histograms show the distribution of the overlap with each randomized eY1H network, where the eY1H network was randomized 10,000 times by edge-switching. The numbers under the histograms indicate the average overlap in 10,000 randomized networks, while the red arrows indicate the observed overlap with the actual eY1H-derived network. Statistical significance was calculated from Z-score values assuming normal distribution for overlap with the randomized networks. (I) Odds ratio for TFs in the eY1H cytokine network having an association with immune diseases in humans (GWAS) or immune phenotypes in KO mice (MGI). Statistical significance determined by Chi-square tests. (J) Schematic of luciferase reporter assays. HEK 293T are co-transfected with TFs fused to the VP160 (10 copies of VP16) activation domain and cytokine promoters (DNA-baits) cloned upstream of a luciferase reporter gene. After 48 h, cells are harvested and luciferase assays performed to measure the level of luciferase light signal. Each PDI is tested in triplicate. (K) Volcano plot of luciferase reporter assays results. Each point represents the average luciferase light signal for each PDI tested in triplicate relative to cells co-transfected with the empty VP160-vector control. PDIs with a fold-change above 1.5 and an FDR-corrected P-value below 0.05 are considered positive PDIs. PDIs that did not pass the selected threshold are also shown. The positive control set consists of PDIs found in the literature or ChIP-seq, and the negative control set consists of non-interacting TF-cytokine pairs.

### A comprehensive eY1H-derived cytokine PDI network

To delineate a comprehensive PDI network for cytokine genes, we used eY1H assays to systematically test 121,632 pairwise interactions between 1,086 human TFs (66% of human TFs) and 112 cytokine promoters (83% of cytokine genes) (**Figure 1C**). The resulting eY1H-derived network comprises 1,380 PDIs between 265 TFs and 108 cytokine promoters (**Figure 1D** and **Table S3**). All the interactions from the eY1H- and literature-derived networks can be searched and visualized in CytReg v2 (https://cytreg.bu.edu/search_v2.html). The eY1H data substantially expands the current literature-derived cytokine PDI network by increasing the number of PDIs, TFs, and cytokines genes. In total, we found 116 TFs that were not previously reported in the literature or by ChIP-seq to bind to or regulate cytokine transcriptional control regions. Of these, 31 TFs have previously been associated with immune diseases in GWAS and/or immune phenotypes in KO mouse studies (MGI). In addition, we identified PDIs for 32 human cytokine genes missing from the literature-derived network, expanding the number of cytokine genes with at least one PDI to 129 (out of ∼140).

We also found that the eY1H-derived network is less biased towards highly studied TFs and cytokines compared to the literature-derived network (**Figure 1E**). Importantly, we observed a similar distribution between the fraction of eY1H-derived PDIs, the expression of TFs in immune cells, and the association of TFs with immune diseases determined by GWAS, across the TF bins (**Figure 1B** and **Table S2**). The association of TFs with immune phenotypes reported from KO mouse studies (MGI) shows an intermediate distribution pattern between the number of eY1H- and literature-derived PDIs, which is unsurprising given that the data in MGI was manually curated from the literature and would be subject to similar biases as the literature-derived PDIs. Overall, the eY1H network expands the cytokine gene PDI network by capturing a broader spectrum of TFs that bind to cytokine promoters and by providing coverage for lesser studied TFs and cytokine genes.

To assess the quality of the eY1H network generated, we compared the eY1H PDIs to interactions identified or predicted by other methods. First, we compared the overlap with literature-derived PDIs and found that among the most highly studied TFs and cytokines, where the literature-derived network is most complete, the overlap between the eY1H- and literature-derived networks is highest, with 44% of the PDIs found by eY1H assays being previously reported (**Figure 1D** and **1F**). Second, we found a significant overlap with ChIP-seq interactions obtained from the ENCODE Project (Encode Project Consortium, 2012) and the GTRD database (Yevshin et al., 2017) (**Figure 1G**), showing that eY1H PDIs recover physical interactions found *in vivo*. This overlap is higher for highly studied TFs, which have been more frequently tested by ChIP-seq (**Figure 1D**). Third, we observed a significant overlap with PDIs predicted based on motif analyses (**Figure 1H**), indicating that the eY1H PDIs are likely a result of direct DNA binding. Finally, we found that TFs in the eY1H-derived network are more frequently associated with immune disorders in humans (GWAS) and immune phenotypes in KO mice (MGI) compared to TFs that did not bind to the cytokine promoters in eY1H assays (**Figure 1I** and **Table S2**), providing confidence that the TFs and PDIs identified by eY1H assays are immune-relevant.

In addition to the *in silico* validation, we experimentally evaluated the quality of the eY1H PDIs by performing luciferase reporter assays in human cells. We tested the ability of TFs fused to the VP160 (10 copies of VP16) activation domain to bind to their target cytokine promoters in HEK 293T cells (**Figure 1J**). We randomly tested 241 eY1H interacting pairs and, at a threshold of >1.5 fold-change (FC) with an FDR < 0.05, we observed a 63% validation rate (**Figure 1K**). Among the eY1H PDIs, we tested 61 PDIs that were previously reported in the literature (22 PDIs) and/or identified by ChIP-seq (43 PDIs) and observed a 72% validation rate, similar to the validation rate of eY1H PDIs for which experimental evidence was not available. As a negative control, we also tested 20 non-interacting TF-cytokine pairs absent from the eY1H-, literature-, or ChIP-seq-derived networks, of which none passed the selected threshold. Overall, the similarly high validation rates of the eY1H and the reported PDIs confirms the high quality of the eY1H PDI dataset.

### Expansion of the cytokine PDI network

The eY1H network includes 37 PDIs that have been reported in the literature, such as interactions between TFAP2A and the promoters of *CXCL8* and *TNF*, and between SPI1 and the *IL12B* promoter (**Table S1**). Additionally, we provide physical binding evidence for 22 literature PDIs that were reported only from functional assays, such as interactions between TFAP2B and the *TNF* gene, and between MAFB and the *IL10* gene (**Table S1**). We also detected 18 PDIs that have been reported in mice but not yet in humans, including interactions between ETS1 and the *CCL5* promoter, and between SMAD4 and the promoters of *CCL2, IFNB1, IL5, IL9*, and *IL10*, showing that these PDIs are conserved between mouse and human.

We found that paralogous and functionally related cytokines often share TF interactors. For example, we found that PPARA, which was reported to regulate the expression of *IL17A* (**Table S1**), also binds to the promoter of *IL17F* (**Figures 2A-B**). Similarly, we detected multiple TF interactors with the promoter of *IL25*, an IL17 family member absent from the literature-derived network, and these TF interactors are shared with *IL17A* and *IL17F*: RORA, RORC, PPARG, NR2F6, and ZNF281 (**Figure 2A**). Indeed, we confirmed that RORC and PPARG bind the *IL25* promoter using luciferase assays in HEK 293T cells (**Figure 2B**). Additionally, we found novel TF interactors shared by *IL17A, IL17F*, and *IL25*, including the nuclear receptors NR1H4, NR2C2, and RXRG (**Figure 2A**). Interestingly, the NR1H4 antagonist ursodeoxycholic acid was reported to inhibit IL17A production and attenuate rheumatoid arthritis, although through a different mechanism (Lee et al., 2017), and RXR agonists were found to suppress *Il17a* expression in mouse (Takeuchi et al., 2013). Altogether, these observations show that in some cases, such as the IL17 family, functionally related cytokines are regulated by shared transcriptional networks.

**Figure 2.**
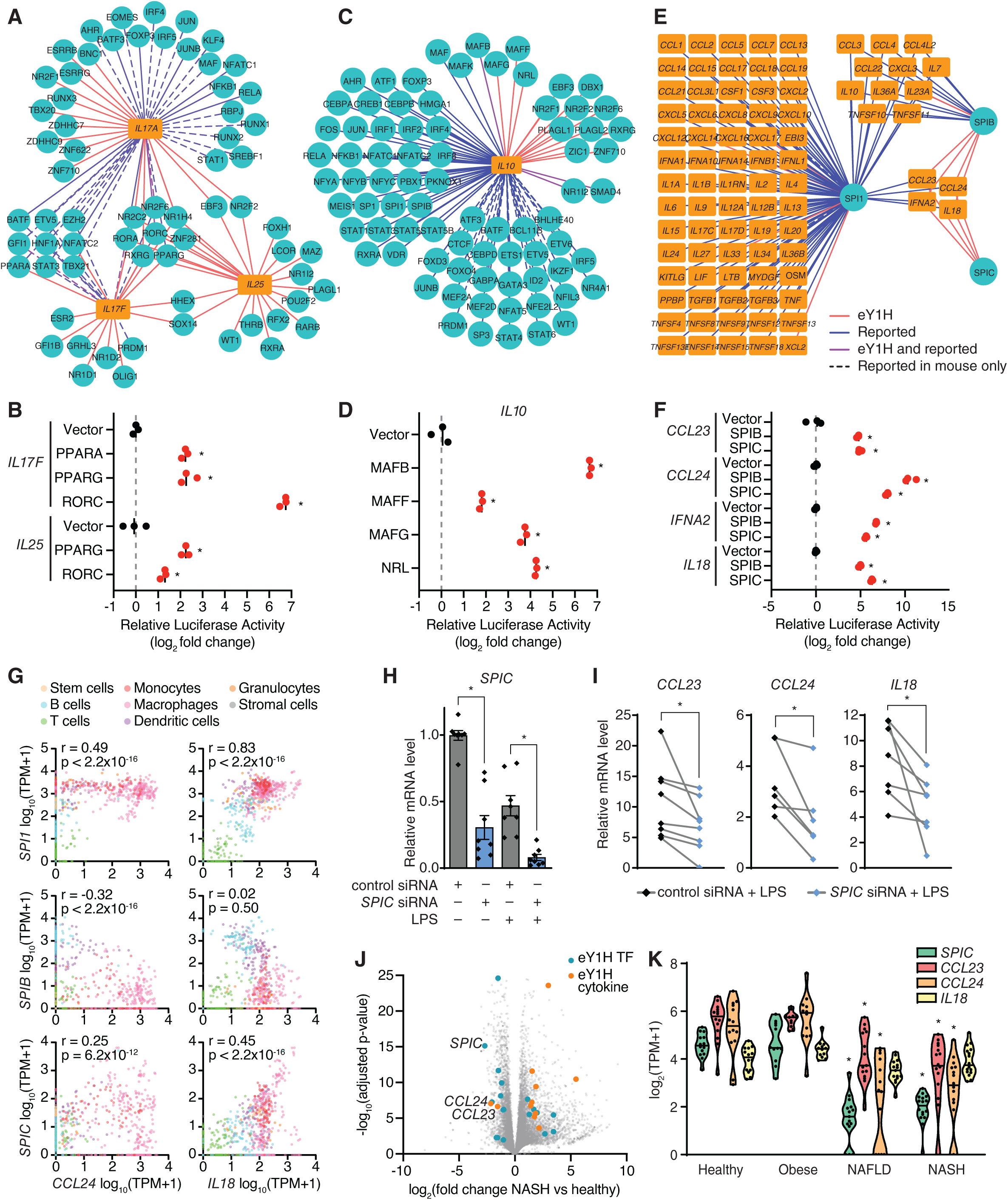
Expansion of the cytokine PDI network. (A, C, and E) eY1H interactions involving IL17 cytokines (A), *IL10* (C), and SPI TFs (E). TFs (turquoise circles) and cytokines (orange rectangles) are connected by edges based on PDIs identified in eY1H assays (red lines), literature or ChIP-seq (blue), or both (purple). PDIs reported in mice but not yet in humans are indicated by dashed lines. (B, D, and F) Luciferase assays performed in HEK 293T cells to validate eY1H interactions. TFs fused to the VP160 activation domain were co-transfected with cytokine promoters cloned upstream of a luciferase gene. The relative luciferase activity is plotted as a fold-change compared to cells co-transfected with the vector control. Experiments were performed in triplicates and the average luciferase activity is indicated by the black line. *P<0.05 by unpaired two-tailed Student’s t-test. (G) Correlations between the expression of *Spi* TFs and *Ccl24* or *Il18* in various immune cells reported in ImmGen. Correlations were determined by Pearson correlation coefficient. (H) Expression of *SPIC* in PBMC-derived macrophages transfected with control non-targeting siRNA or siRNA targeting *SPIC*, and stimulated with 10 ng/ml LPS for 4 hours. Error bars indicate the standard error of the mean. *P<0.05 by paired Wilcoxon signed-rank test. (I) Expression of SPIC target cytokines (*CCL23, CCL24*, and *IL18*) in PBMC-derived macrophages transfected with control non-targeting siRNA or siRNA targeting *SPIC*, and stimulated with 10 ng/ml LPS for 4 hours. Each line represents a different donor evaluated in biological triplicate. *P<0.05 by paired Wilcoxon signed-rank test. (J) Volcano plot of differential gene expression between healthy and non-alcoholic steatohepatitis (NASH) liver biopsies. eY1H TFs (blue) and cytokines (orange) with FC ≥ 2.0 and adjusted p-value <0.05 are indicated. (K) Expression of *SPIC* and its target cytokines (*CCL23, CCL24*, and *IL18*) in liver biopsies from healthy, obese, non-alcoholic fatty liver disease (NAFLD), and NASH patients. *P<0.05 by Kruskal-Wallis test with Dunn’s multiple comparisons test when compared to healthy patients.

We also observed that TFs from the same family often interact with the same cytokine promoters. Although this may be due to similar DNA-binding specificities shared between TF paralogs, it may also be associated with TF redundancy, division of labor between TFs expressed in different cells or different biological contexts, and TFs with opposing functions as we have previously shown for TFs regulating developmental enhancers (Fuxman Bass et al., 2015). The Maf TFs, MAF, MAFB, and MAFK, have been shown to regulate *IL10* (**Table S1**). Using eY1H and luciferase assays we detected interactions between their paralogs, MAFF, MAFG, and NRL, and the *IL10* promoter (**Figures 2C-D**). Similarly, we found that SPIB and SPIC bind to the promoters of four target cytokine genes, *CCL23, CCL24, IFNA2*, and *IL18* (**Figure 2E**), which were previously found by ChIP-seq to be bound by their close paralog SPI1 (also known as PU.1) (Encode Project Consortium, 2012). We further confirmed these PDIs by luciferase assays in HEK 293T cells (**Figure 2F**). Interestingly, SPI1, SPIB, and SPIC have been reported to have redundant and opposing functions in B cell development (Li et al., 2015; Schweitzer et al., 2006; Willis et al., 2017).

In mouse immune cells reported by the Immunological Genome Project (ImmGen) (Yoshida et al., 2019), the expression of *Spi1* and *Spic* are positively correlated, while the expression of *Spib* is generally negatively correlated with their target cytokines, *Ccl24* and *Il18* (**Figure 2G**). To our knowledge, SPIC has not yet been reported to directly regulate the expression of cytokines genes. When we knocked down the expression of *SPIC* by siRNA in human primary macrophages (**Figure 2H**), the LPS-induced expressions of *CCL23, CCL24*, and *IL18* were also downregulated (**Figure 2I**), suggesting that SPIC activates the expression of these cytokine genes. *Spic* is highly expressed in lymphocytes and lymphoid tissues, where it has been reported to regulate the development of macrophage populations (Haldar et al., 2014; Kohyama et al., 2009). *SPIC* is also expressed in the liver (Human Protein Atlas, (Uhlen et al., 2015)), possibly in Kupffer cells. In liver biopsies from patients diagnosed with non-alcoholic fatty liver disease (NAFLD) and its inflammatory subtype non-alcoholic steatohepatitis (NASH), in which Kupffer cells and recruited macrophages play a central role in disease progression (Kazankov et al., 2019), the expression of *SPIC* and its target cytokines, *CCL23* and *CCL24*, were significantly downregulated (**Figures J-K**) (Suppli et al., 2019). Given the role of CCL23 in monocyte chemotaxis and angiogenesis, and the role of CCL24 in eosinophil recruitment, all of which contribute to liver inflammation in NASH (Forssmann et al., 1997; Hart et al., 2017; Hwang et al., 2005), *SPIC* downregulation may constitute a compensatory homeostatic mechanism to limit disease pathogenesis.

### Role of nuclear receptors in modulating the expression of cytokine genes

We detected multiple TF families in the eY1H-derived cytokine network, including major families such as Cys2His2 zinc fingers (ZF-C2H2) and homeodomains (**Figure 3A** and **Table S3**). We found an enrichment of interactions with the AP-2 and IPT/TIG families (p = 1.0 x 10^−9^ and p = 6.2 x 10^−8^ by proportion comparison test, respectively), which are known to play prominent roles in immune cell differentiation and immune responses (Rao et al., 1997). Interestingly, we also observed a significant enrichment of PDIs involving nuclear receptors (NRs) (p = 7.4 x 10^−28^ by proportion comparison test). While NRs represent <4% of the 1,086 TFs tested by eY1H assays, they constitute 18% of the PDIs (248 PDIs involving 28 NRs) in the eY1H network (**Figures 3A-B**). This also represents a 3.5-fold increase in the number of PDIs involving NRs compared to PDIs reported in the literature (**Table S1**).

**Figure 3.**
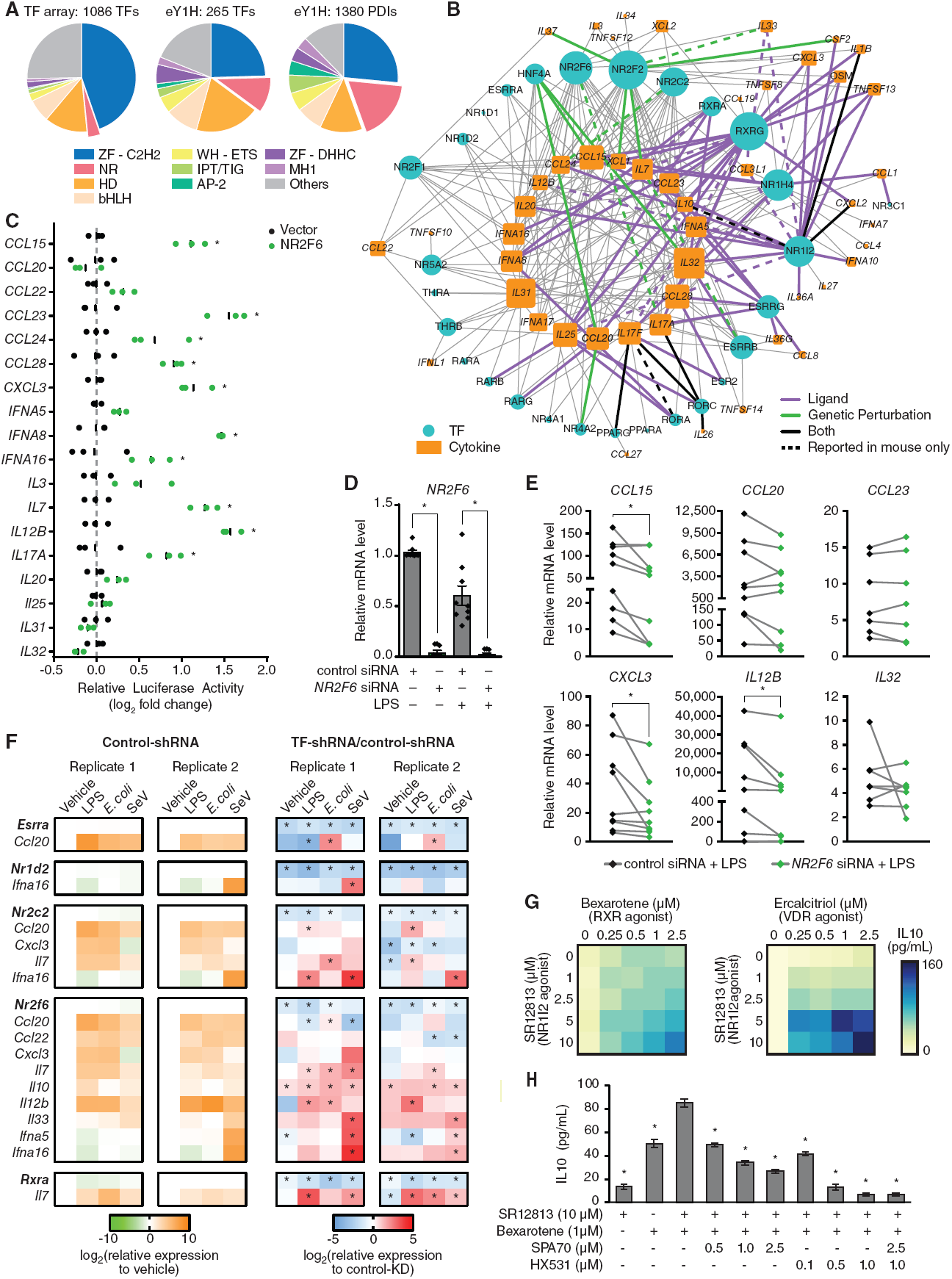
Role of nuclear receptors in modulating the expression of cytokine genes. (A) Pie charts showing the distribution of TF families in the 1,086 TFs tested by eY1H assays and the TFs and PDIs in the eY1H-derived network. AP2 – activating protein 2; bHLH – basic helix-loop-helix; HD – homeodomain; IPT/TIG – immunoglobulin-like, plexins, TFs/TF immunoglobulin; MH1 – Mad homology 1; NR – nuclear receptor; WH-ETS – winged-helix E-twenty six; ZF-C2H2 – zinc finger C2H2; ZF-DHHC – zinc finger DHHC. (B) eY1H-derived PDI network involving nuclear receptors (NRs) comprising 248 PDIs (edges) between 28 NRs (turquoise circles) and 48 cytokine promoters (orange rectangles). PDIs reported in the literature or NURSA from functional assays using ligands (purple edges), genetic perturbation studies (green edges), or both (black edges) are indicated. PDIs determined in mice but not yet in humans are indicated by dashed lines. The size of the node indicates the number of interactors. (C) Luciferase assays performed in HEK 293T cells co-transfected with an NR2F6 expressing vector and 18 cytokine promoters cloned upstream of a luciferase gene. The relative luciferase activity is plotted as a fold-change compared to cells co-transfected with the vector control. Experiments were performed in triplicate and the average luciferase activity is indicated by the black line. *P<0.05 by unpaired two-tailed Student’s t-test and FC>1.5. (D) Expression of *NR2F6* in human PBMC-derived macrophages transfected with control non-targeting siRNA or siRNA targeting *NR2F6*, and stimulated with 10 ng/ml LPS for 4 hours. Error bars indicate the standard error of the mean. *P<0.05 by paired Wilcoxon signed-rank test. (E) Expression of NR2F6 target cytokines in human PBMC-derived macrophages transfected with control non-targeting siRNA or siRNA targeting *NR2F6*, and stimulated with 10 ng/ml LPS for 4 hours. Each line represents a different donor evaluated in biological triplicate. *P<0.05 by paired Wilcoxon signed-rank test. (F) Expression of TFs and cytokines in mouse iBMDMs transduced with control luciferase-targeting shRNA or shRNA targeting mouse nuclear receptors (*Esrra, Nr1d2, Nr2c2, Nr2f6, or Rxra*), and stimulated with 1 µg/ml LPS, 1 CFU *E. coli*, or 200 HAU Sendai virus. The log_2_ fold-change expression of TFs and cytokines in response to the ligands versus vehicle in control-shRNA iBMDMs are shown on the left (green-orange heatmaps). The log_2_ fold-change in relative expression of TFs and cytokines in response to the ligands between TF-shRNA and control-shRNA iBMDMs are shown on the right (blue-red heatmaps). The experiment was performed twice and each condition was tested in biological triplicate. *P<0.05 by unpaired two-tailed Student’s t-test. (G) Heatmap showing the expression of IL10 (pg/ml) in THP-1-derived M2 macrophages treated with increasing concentrations of SR12813 (NR1I2 agonist) and Bexarotene (RXR agonist) or Ercalcitriol (VDR agonist). Experiments were performed in quadruplicate and the average expression of IL10 (pg/ml) is shown. Data is representative of 2 experiments. (H) Expression of IL10 (pg/ml) in THP-1-derived M2 macrophages treated with SR12813 (NR1I2 agonist) and Bexarotene (RXR agonist), SPA70 (NR1I2 antagonist), and/or HX531 (RXR antagonist). Experiments were performed in quadruplicate and the average expression of IL10 (pg/ml) is shown. Error bars indicate the standard error of the mean. *P<0.05 by unpaired two-tailed Student’s t-test when compared to THP-1-derived M2 macrophages treated with 10 µM SR12813 + 1 µM HX531.

NRs are a family of 48 ligand-activated TFs (Zhang et al., 2004) that can sense multiple endogenous and exogenous ligands (*e.g.*, steroids, retinoids, vitamins, xenobiotics, and other lipophilic substances) and modulate the expression of genes involved in a variety of biological processes including development, differentiation, metabolism, and immunity (Sever and Glass, 2013). Indeed, NRs have been shown to modulate the expression of key cytokines, such as *Il17a* (Hermann-Kleiter et al., 2012; Ivanov et al., 2006) and *TNF* (Jiang et al., 1998), in immune cell differentiation and autoimmune disorders. To determine whether NRs identified by eY1H assays potentially regulate their target cytokine genes, we searched for functional evidence for the eY1H PDIs in the literature and in expression profiling datasets from the Nuclear Receptor Signaling Atlas (NURSA) Transcriptomine database (Becnel et al., 2017). Notably, for 80 of the 248 PDIs, the NR has been found to functionally regulate its target cytokine gene (**Figure 3B** and **Table S4**); however, in most cases, direct DNA binding had not been reported. The majority of reported regulatory interactions involving NRs have only been studied using natural or synthetic ligands. For example, NR1I2 (also known as PXR) is activated by a variety of compounds including steroids, antibiotics, bile acids, and plant metabolites (Willson and Kliewer, 2002), and 18 of the 20 cytokine genes found to interact with NR1I2 by eY1H assays were reported to be functionally regulated by NR1I2 ligands (**Figure 3B** and **Table S4)**. NR1I2 has been highly studied for its role in metabolism, but has also recently emerged as an important regulator of innate immune responses (Wang et al., 2014), T-cell function (Dubrac et al., 2010), and inflammatory bowel disease (IBD) (Cheng et al., 2012). Similarly, 19 of the 29 cytokine genes found to interact with RXRG were reported to be regulated by RXR ligands such as vitamin A derivatives and fatty acids (**Figure 3B** and **Supplementary Table S4)**. RXRA is well known to regulate inflammatory responses and attenuate host antiviral responses by modulating cytokine expression (Ma et al., 2014; Nunez et al., 2010), and RXRA deficiency has been associated with autoimmune diseases (Roszer et al., 2011). Less is known about its paralog RXRG, for which we found many known functional and novel interactions by eY1H assays (**Figure 3B** and **Table S4**).

Some NRs, such as NR2C2, NR2F2, and NR2F6, have no known natural or synthetic ligand, but have been studied using genetic perturbations. For example, ChIP-seq and microarray analysis of *NR2F2* knockdown in endometrial stromal cells show that NR2F2 primarily regulates the expression of genes involved in inflammation and cytokine signaling (Li et al., 2013). Its paralog, NR2F6, has been reported to be a critical regulator of Th17 cell fate and function by binding to the *Il17a* promoter, thereby preventing the binding of NFAT/AP-1 and RORC (Hermann-Kleiter et al., 2012). To determine whether NR2F6 can functionally regulate the promoters of other cytokine genes found by eY1H assays, we performed luciferase assays in HEK 293T cells and found that NR2F6 activated 10 of the 18 cytokine promoters tested (**Figure 3C**). Additionally, we found that in human primary macrophages in which we knocked down the expression of *NR2F6* by siRNA (**Figure 3D**), the LPS-induced expressions of *CCL15, CXCL3*, and *IL12B*, were also downregulated (**Figure 3E**). Taken together, these observations show that, contrary to the mechanism described in mouse Th17 cells, NR2F6 activates the expression of cytokine genes in human macrophages.

Cytokines are often stimulated in response to pathogen-associated molecular patterns (e.g., LPS, CpG DNA, and peptidoglycan) and pathogens. To determine whether other NRs in the eY1H network modulates cytokine expression in stimulated conditions, we knocked down five NRs in mouse immortalized bone marrow-derived macrophages (iBMDMs) using a constitutive shRNA system and treated the iBMDMs with LPS, *E. coli*, or Sendai virus (**Figure 3F**). Of the 16 TF-cytokine pairs tested, 15 resulted in the modulation of cytokine expression induced by at least one of the stimuli in either replicate, and 10 were significant for the same stimuli in both replicates. Some TFs showed stimuli-specific regulation of their target cytokines. For example, *Esrra* knockdown significantly increased the expression of *Ccl20* only in *E. coli-*stimulated conditions, and *Nr2c2* knockdown significantly increased the expression of *Ccl20* only in LPS-stimulated conditions. Other TFs showed more general regulatory mechanisms that may be condition-independent, such as *Nr2f6* knockdown that resulted in increased expression of *Il7* and *Il10* in all conditions tested. Interestingly, while NR2F6 activated cytokine genes in human cells (**Figure 3C** and **3E**), consistent with observations in mouse Th17 cells (Hermann-Kleiter et al., 2012), NR2F6 repressed cytokine gene expression in mouse iBMDMs (**Figure 3F**), suggesting a different regulatory mechanism between human and mouse. Overall, these observations show that NRs can modulate the expression of cytokine genes, and that some of these regulatory interactions occur only under specific stimulated conditions.

### Modulation of cytokine expression using synergistic TF combinations

Modulation of cytokine expression using small molecules or blocking cytokine activity using antibodies have been used as potent therapeutic strategies for multiple immune-related diseases (Schett et al., 2013). NRs present promising therapeutic drug targets to modulate cytokine expression because of the lipophilic nature of their ligands and because multiple NR agonists and antagonists have been approved as therapeutic agents (Sonoda et al., 2008). To explore the therapeutic potential of NRs in modulating cytokine expression, we focused on IL10, an important anti-inflammatory cytokine that is often downregulated in autoimmune diseases such as IBD (Neumann et al., 2019). M2 macrophages are a major source of IL10 (Roszer, 2015); however, THP-1-derived M2 macrophages do not readily produce IL10 (Schildberger et al., 2013) and thus serve as a fitting model to test for pairs of NRs that synergistically promote IL10 production. From eY1H assays, we found several NRs including NR1I2 and RXRG that bind to the *IL10* promoter, while RXRA and VDR have been reported to regulate *IL10* (**Table S1**). Strikingly, we found that in THP-1-derived M2 macrophages, combinations of SR12813 (NR1I2 agonist) with Bexarotene (RXRA and RXRG agonist) or with Ercalcitriol (VDR agonist) synergistically promoted IL10 production (**Figure 3G**). Further, SPA70 (NR1I2 antagonist) and HX531 (RXRA and RXRG antagonist) reduced the IL10 production mediated by SR12813 and Bexarotene (**Figure 3H**). Altogether, these observations demonstrate the use of the eY1H- and literature-derived networks as a framework to identify combinations of TFs that synergistically regulate the expression of target cytokine genes.

### Prediction of disease-associated PDIs

Cytokine imbalance is widely associated with the pathogenesis of immune disorders including autoimmunity and susceptibility to infections (Turner et al., 2014). Similarly, aberrant TF expression or activity is also associated with immune disorders, likely due to dysregulation of downstream target genes important for immune cell functions such as cytokines genes (Barnes and Karin, 1997). Indeed, we found many TFs in the eY1H network that have been associated with the same immune disorders (reported in DisGeNET, GWAS, or MGI) as their corresponding target cytokines (**Figure 4A**). In total, we found 377 TF-cytokine-disease associations, involving 73 TFs and 60 cytokines, wherein both the TF and target cytokine have been associated with the same immune disease (**Figure 4A, Figure S1**, and **Table S5**), which is higher than expected by chance (**Figure 4B**). We found known TF-cytokine disease regulatory axes, such as RORC inducing IL17A and IL17F in inflammatory arthritis, psoriasis, and multiple sclerosis (Ivanov et al., 2006; Venken et al., 2019). In other cases, the link is less well-established. For example, loss of GFI1 results in de-repression of *Il17f* in type 2 innate lymphoid cells (ILC2) in the course of infection (Spooner et al., 2013). However, evidence of the GFI1-IL17F regulatory axis in asthma has not been determined. For most TF-cytokine-disease associations, the link has not yet been explored, and thus, the identified associations constitute a framework to generate hypotheses of regulatory axes in disease contexts.

**Figure 4.**
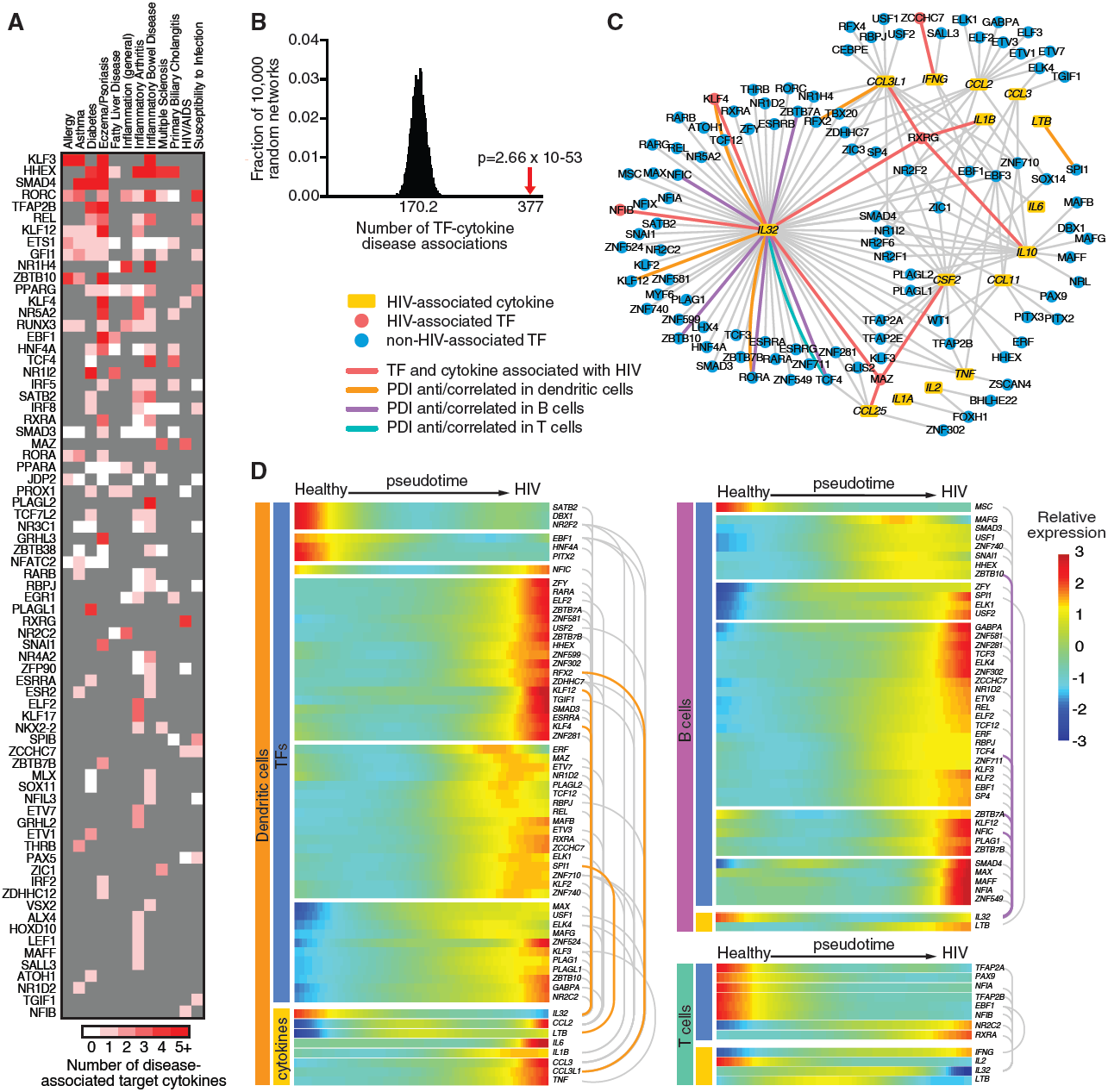
HIV-associated PDI network. (A) Heatmap showing TFs and number of target cytokines (shades of red) associated with the same immune disease (reported in DisGeNET, GWAS, or MGI). TFs not associated with the immune disease are shaded in grey. (B) eY1H PDIs include significantly more TF-cytokine-disease associations than expected by chance. The eY1H network was randomized 10,000 times by edge-switching and the overlap of TF-cytokine-disease associations with each randomized network was calculated. The numbers under the histogram indicates the average overlap in 10,000 randomized networks, while the red arrow indicates the observed overlap with the actual eY1H-derived network. Statistical significance was calculated from Z-score values assuming normal distribution for overlap with the randomized networks. (C) HIV-associated PDI network comprising 171 PDIs (edges) between 15 cytokines (yellow rectangle nodes) associated with HIV (reported in DisGeNET, GWAS, or MGI) and 102 TF interactors (circle nodes). TFs associated with HIV are represented by red nodes, while TFs not yet associated with HIV are represented by blue nodes. PDIs between TFs and cytokines that are both associated with HIV are indicated by red edges. PDIs correlated/anti-correlated in expression on a single-cell basis in dendritic cells (orange edges), B cells (purple edges), or T cells (green edges), are indicated. (D) TFs and cytokines in the HIV-associated PDI network that significantly change in expression along the pseudotime disease trajectory in each cell type. eY1H PDIs are connected by grey arches, and PDIs correlated/anti-correlated in expression on a single-cell basis in dendritic cells (orange arches), B cells (purple arches), or T cells (green arches), are indicated.

### HIV cytokine PDI network

To find supporting evidence for our predicted TF-cytokine-disease associations, we integrated scRNA-seq expression data to reveal correlated TF-cytokine pairs at the single-cell level that may contribute to pathogenesis of the disease. As a proof-of-concept, we focused on HIV given its importance in human health and the availability of scRNA-seq data from healthy and infected individuals (Farhadian et al., 2018). The HIV-associated PDI network contains 171 PDIs between 15 cytokines associated with HIV (reported in DisGeNET, GWAS, or MGI) and 102 TF interactors (**Figure 4C** and **Table S5**). Indeed, we found ten PDIs wherein the cytokine and its TF interactor were both found to be associated with HIV. To identify correlated PDIs, we integrated scRNA-seq data collected from cerebrospinal fluid and PBMC-derived dendritic cells, B cells, and T cells, from HIV-infected individuals (Farhadian et al., 2018) (**Figure S2A-B)**. In total, we identified nine TF-cytokine pairs correlated/anti-correlated in expression on a single-cell basis in at least one cell type, of which two pairs were correlated/anti-correlated in multiple cell types from HIV-infected individuals (**Figure 4C** and **Table S6**). For example, we found that *IL32*, a cytokine shown to suppress HIV replication (Rasool et al., 2008), is anti-correlated with *KLF4* and *KLF12* expression on a single-cell basis in dendritic cells from HIV-infected individuals (**Figure 4C** and **Table S6)**, suggesting that the KLF4 and KLF12 TFs may repress *IL32* expression in HIV. While KLF4 has been associated with HIV, KLF12 has not. Interestingly, the *KLF12* gene region was found to exhibit lower methylation levels and was enriched in H3K27ac histone modifications in HIV-infected subjects compared to uninfected subjects (Zhang et al., 2016), suggesting that KLF12 dysregulation may contribute to *IL32* repression in HIV infection.

To determine whether correlations at the single-cell level could be related to the course of HIV progression, we used the scRNA-seq data to reconstruct a pseudotime disease trajectory. Briefly, a differential expression analysis was performed to identify the top significantly differentially expressed genes between healthy and HIV-infected conditions to build the disease trajectory, and then each single-cell was assigned a numeric pseudotime value and then ordered along the disease trajectory (**Figure S2C-E**). Across dendritic cells, B cells, and T cells, we found that 67 of 102 TFs, which are involved in 69 PDIs with HIV-associated cytokines, significantly changed in expression along the pseudotime trajectory in at least one cell type (**Figure 4D**). For example, *KLF4, KLF12*, and *IL32* were found to significantly change in expression along the pseudotime and both *KLF4-IL32* and *KLF12-IL32* TF-cytokine pairs were found to be anti-correlated along the pseudotime in dendritic cells (**Figure 4D)**. Although some of the TFs identified may change in expression as a consequence of HIV disease progression, others may lead to changes in expression of their target cytokines, even if a correlation at the single-cell level was not detected (*e.g.*, due to changes in TF activation rather than expression).

### Inflammatory bowel disease cytokine PDI network

Using a similar approach, we next focused on IBD given the important role of cytokines in IBD pathogenesis (Neurath, 2014). The IBD-associated PDI network contains 582 PDIs between 46 cytokines associated with IBD (reported in DisGeNET, GWAS, or MGI) and 195 TF interactors (**Figure 5A**). Importantly, we found 71 PDIs wherein the cytokine and its TF interactor were both found to be associated with IBD. To identify correlated TF-cytokine pairs involved in PDIs, we integrated scRNA-seq data collected from intestinal epithelial cells, dendritic cells, macrophages, B cells, and T cells, from IBD patients (Martin et al., 2019) (**Figure S3)**, and found 62 TF-cytokine pairs correlated/anti-correlated in expression on a single-cell basis in at least one cell type, of which 12 PDIs were correlated/anti-correlated in multiple cell types (**Figure 5A** and **Table S7)**. Further, 18 PDIs correlated/anti-correlated in expression involve TFs that are associated with IBD, such as RORC, RUNX3, and NR1H4, which have been shown to play a role in IBD by directly modulating the expression of cytokine genes (Brenner et al., 2004; Gadaleta et al., 2011; Sawa et al., 2011). Notably, nine PDIs correlated/anti-correlated in expression have been reported in the literature. Five of these PDIs involve TFs that are associated with IBD (i.e., REL-*CXCL8*, REL-*CCL20*, RORC-*IL17A*, RORC-*IL26*, and NR4A2-*CCL20*), suggesting these TF-cytokine pairs may constitute disease regulatory axes in IBD. The other four PDIs involve TFs that are not yet associated with IBD (*i.e.*, RORA-*IL17F*, MAFB-*IL10*, TFAP2A-*CXCL8*, and ELK1-*CCL2*), and further studies are needed to determine their role in IBD. Taken together, these findings show that the IBD-associated PDI network provides a framework to identify TF-cytokine regulatory axes that contribute to pathogenesis of the disease.

**Figure 5.**
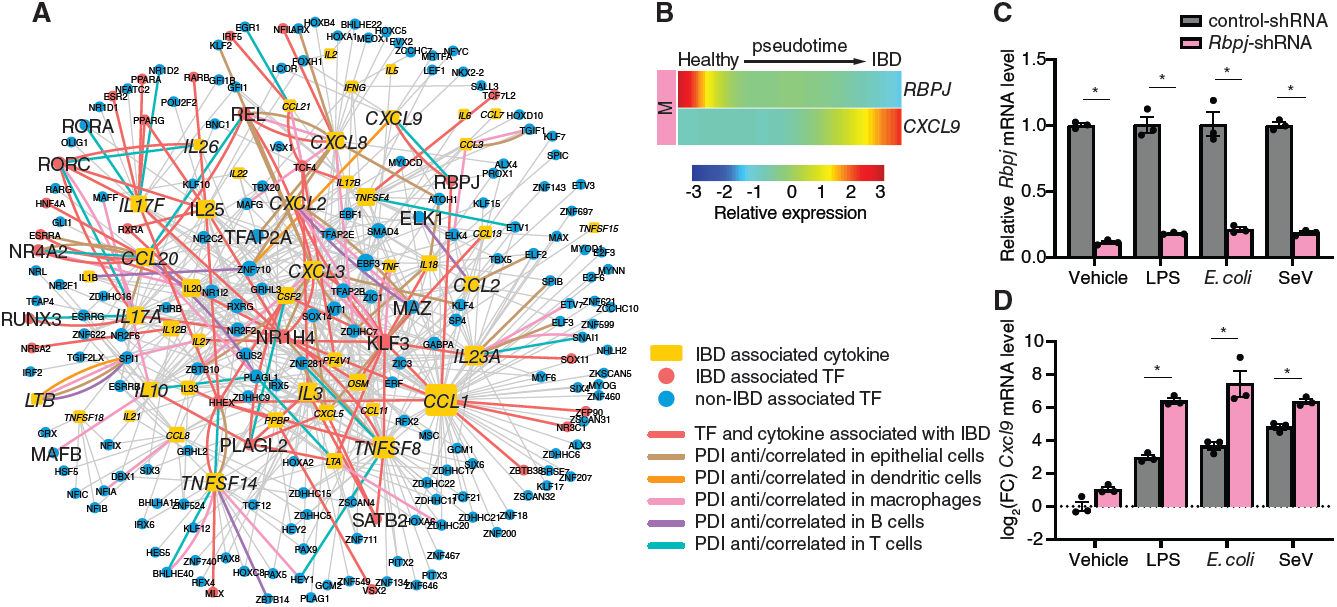
IBD-associated PDI network. (A) IBD-associated PDI network comprising 582 PDIs (edges) between 46 cytokines (yellow rectangle nodes) associated with IBD (reported in DisGeNET, GWAS, or MGI) and 195 TF interactors (circle nodes). TFs associated with IBD are represented by red nodes, while TFs not yet associated with IBD are represented by blue nodes. PDIs between TFs and cytokines that are both associated with HIV are indicated by red edges. PDIs correlated/anti-correlated in expression on a single-cell basis in intestinal epithelial cells (brown edges), dendritic cells (orange edges), macrophages (pink edges), B cells (purple edges), or T cells (green edges), are indicated. (B) Expression of *RBPJ* and *CXCL9* along the pseudotime in macrophages are anti-correlated. (C and D) Relative expression of *Rbpj* (C) or log_2_ fold-change expression of *Cxcl9* (D) in mouse iBMDMs transduced with control luciferase-targeting shRNA or shRNA targeting mouse *Rbpj*, and stimulated with 1 µg/ml LPS, 1 CFU *E. coli*, or 200 HAU Sendai virus. Experiments were performed in triplicate and the average expression of *Rbpj* or *Cxcl9* is shown. Error bars indicate the standard error of the mean. Data are representative of 2 experiments. *P<0.05 by unpaired two-tailed Student’s t-test.

To determine whether TF-cytokine pairs in the IBD-associated PDI network could have a role in IBD progression, we used the scRNA-seq data from healthy and IBD patients to reconstruct a pseudotime trajectory of disease progression (**Figure S3C-G**). Across intestinal epithelial cells, dendritic cells, macrophages, B cells, and T cells, we found 141 of 195 TFs and 37 of 46 cytokines that significantly changed in expression along the pseudotime trajectory in at least one cell type (**Figure S4**). To identify novel TF-cytokine disease regulatory axes, we focused on 42 PDIs between TFs and cytokines that are both associated with IBD and significantly change in expression along the pseudotime. Interestingly, *RBPJ* and *CXCL9* were found to be anti-correlated along the pseudotime in macrophages (**Figure 5B**). In PBMC-derived monocytes infected with Salmonella, a bacterial disease that affects the intestinal tract, *RBPJ* and *CXCL9* were also anti-correlated in expression on a single-cell basis (Bossel Ben-Moshe et al., 2019). Further, mice harboring a *RBPJ* deletion spontaneously develop IBD (Obata et al., 2012), while experimental IBD models and IBD patients manifest significantly increased *CXCL9* levels (Lacher et al., 2007; Singh et al., 2007). To determine whether RBPJ regulates *CXCL9* in infection or inflammation, we knocked down *RBPJ* in iBMDMs and treated the iBMDMs with LPS, *E. coli*, or Sendai virus (**Figure 5C**). *RBPJ* knockdown significantly increased the expression of *CXCL9* across all conditions (**Figure 5D**), providing evidence of a disease regulatory axis involving a loss of RBPJ-mediated repression of *CXCL9* in infection and inflammation. Altogether, these findings demonstrate the power of the eY1H network to identify novel TFs and PDIs that have a role in inflammatory diseases.

### Identification of lineage TFs regulating cytokine genes

Cytokines have a fundamental role in the determination and commitment to specific immune cell lineages. For example, IFNG promotes Th1-cell differentiation and inhibits Th2-cell differentiation, whereas IL4 promotes Th2-cell differentiation and inhibits Th1-cell differentiation (Ansel et al., 2006). Production of these lineage-directing cytokines is regulated by key transcription factors that ultimately determine the fate of the cell. For example, TBX21 (also known as T-bet) regulates the *IFNG* promoter in Th1 cells and GATA3 regulates the *IL4* promoter in Th2 cells (Ansel et al., 2006). Although cell-fate determining TFs have been identified for many immune cell lineages, TFs involved in early fate decisions and maintenance of the established cell lineages are still being discovered.

To predict novel TFs that have a role in development of immune cell lineages, we leveraged the eY1H network and gene associations with abnormalities in immune cell differentiation (reported in DisGeNET, GWAS, or MGI) to generate lineage-associated PDI networks (**Figure 6**). Each lineage network consists of cytokines that are associated with abnormalities in lineage-specific differentiation and their TF interactors (**Table S8**). Indeed, we found 100 TF-cytokine-lineage associations across all the lineage networks, wherein the cytokine and its TF interactor were both found to be associated with differentiation abnormalities in the same lineage. For example, RORC and IL17A were associated with CD4+ T cell differentiation, SPIC and IL18 were associated with macrophage differentiation, and RUNX3 and XCL1 were associated with dendritic cell differentiation. Further, when we searched for additional TF-lineage associations in the literature, we found an additional 119 TF-cytokine-lineage associations in which the TFs were reported to be associated with differentiation of the same lineage as their target cytokines (**Figure 6** and **Table S9**). We also found several TFs that have not yet been associated but have been predicted to regulate differentiation in the same lineage as their target cytokines. For example, ELF3 was predicted in neutrophil differentiation based on its expression patterns during differentiation (Lee et al., 2015), and PROX1 was predicted to play a role in CD4+ T cell differentiation by suppressing key lineage cytokines (Zhang et al., 2017). Overall, the lineage-associated PDI networks generated from cytokines associated with abnormalities in lineage-specific differentiation, identified many TF interactors found by eY1H assays that have themselves been associated with differentiation in the same lineage.

**Figure 6.**
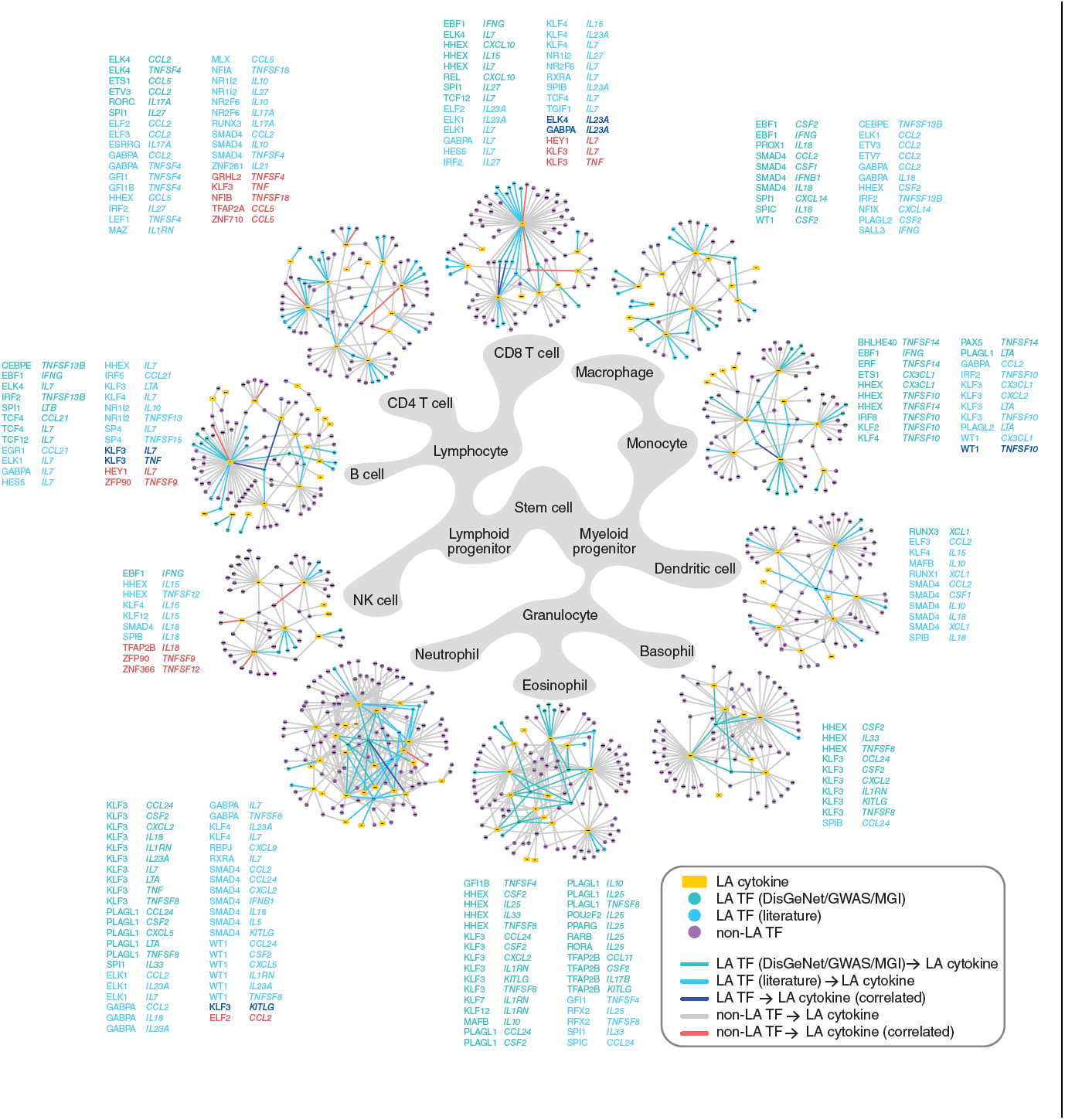
Immune cell lineage PDI networks. Lineage-associated (LA) PDI networks. Each LA network consists of eY1H PDIs involving cytokines (yellow rectangle nodes) associated with lineage-specific differentiation (DisGeNET, GWAS, or MGI), and their TF interactors (circle nodes). TFs associated with abnormalities in lineage-specific differentiation reported in DisGeNET, GWAS, or MGI, are represented by green nodes. TFs associated with lineage-specific differentiation reported in the literature are represented by cyan nodes. TFs not yet associated with lineage-specific differentiation are represented by purple nodes. PDIs between TFs and cytokines both associated with the same lineage-specific differentiation are indicated by green or cyan edges, and listed next to each lineage network in green or cyan font. PDIs between TFs and cytokines found to be correlated by scRNA-seq in the specific lineages are indicated by blue edges for LA TFs, or red edges for non-LA TFs, and listed next to each lineage network in blue or red font, respectively.

To narrow our focus on PDIs likely involved in immune cell differentiation, we integrated scRNA-seq data collected from human hematopoietic stem/progenitor cells (HSPCs) during early cell fate commitment (Pellin et al., 2019). We found 20 PDIs wherein the TF and cytokine are significantly correlated in expression on a single-cell basis in the corresponding cell lineage (**Table S10**). Of these, six PDIs involve TFs that have been associated with lineage differentiation and 14 PDIs involve TFs not yet associated with lineage differentiation. This includes KLF3, which was found to be correlated with TNF in CD4+ and CD8+ T cells. Interestingly, although KLF3 has not yet been associated with T cell differentiation, KLF3 is highly expressed in T cells (ImmGen, (Yoshida et al., 2019)) and is predicted to regulate memory T cell formation (Best et al., 2013). Additionally, some correlated TFs that have not yet been associated with lineage-specific differentiation have been observed to play a role in proliferation. For example, *HEY1*, a target of the NOTCH signaling pathway, was found to be correlated with the essential B cell cytokine IL7. In B cell lymphoma, *HEY1* is overexpressed, and impairing either NOTCH signaling or downregulation of *HEY1* results in reduced proliferation of B cell lymphoma cells (Cao et al., 2014). *IL7* has also been found to be expressed and enhance the development of B cell lymphomas (Rich et al., 1993). These observations suggest that beyond the context of cancer, HEY1 may have a role in B cell development potentially by regulating *IL7*. In total, 26 TFs that have not yet been associated with lineage-specific differentiation have been implicated in proliferative disorders, including neoplastic growth transformations or leukemia, in the same lineage as their target cytokines (**Table S9**), suggesting that these TFs may also have roles in the proliferation stages during lineage development. Altogether, these findings demonstrate the use of the eY1H network to identify novel TFs that have a lineage-defining role in the development of immune cells.

### Supporting evidence for eY1H-derived PDIs

We devised a “supporting evidence score” for each eY1H PDI, in which we weighted the interactions based on: (1) literature-reported evidence, (2) presence of ChIP-seq peaks in the cytokine promoter, (3) presence of corresponding TF-binding motif in the cytokine promoter, (4) functional evidence from genetic or drug perturbation experiments reported in NURSA Transcriptomine, (5) experimental evidence from this project (*e.g.*, luciferase reporter assays and TF knockdown experiments in primary cells), (6) TF-cytokine co-expression in bulk RNA-seq or single-cell RNA-seq datasets analyzed in this project, and (7) shared TF-cytokine associations with immune diseases or lineage differentiation (**Table S11**). In total, 973 PDIs are supported by some evidence of a potential interaction, including 243 PDIs supported by reported experimental evidence (*e.g.*, functional assays, binding assays, or ChIP-seq), and 175 PDIs experimentally validated in this study in mammalian cell lines or primary macrophages. Additionally, 90 PDIs are correlated/anti-correlated in bulk RNA-seq or scRNA-seq datasets suggesting that these PDIs have a regulatory function, and 313 PDIs share the same association with immune diseases or lineage differentiation. Overall, we found abundant evidence supporting the PDIs detected by eY1H assays.

## Discussion

In this project, we used eY1H assays to delineate a comprehensive cytokine PDI network comprising 1,380 PDIs between 265 TFs and 108 cytokine gene promoters. We performed *in silico* and *in vitro* validations confirming the high quality of the network, and further investigated several TFs in primary macrophages showing that they can functionally regulate their target cytokines. Additionally, we found correlations/anti-correlations in expression between many TF-cytokine pairs in bulk or scRNA-seq datasets. Altogether, we found supporting evidence, either from external sources or validations we performed in this study, for over 70% of the PDIs found by eY1H assays. To summarize the supporting evidence, we provide a table (**Table S11**) and an evidence score for each PDI providing a resource for researchers to prioritize PDIs for follow-up studies.

It is important to note that the data used to generate the “supporting evidence score” are not complete and have their own confidence issues. For example, functional validations reported in the literature or phenotypic studies suffer from research biases towards highly studied TFs and cytokines. Additionally, only 30% of human TFs have been tested by ChIP-seq due to the lack of ChIP-grade anti-TF antibodies, and ChIP-seq has only been performed in a limited number of cell types and conditions, mostly non-immune and unstimulated cell lines. Thus, interactions not supported by any evidence may still be physiologically relevant, and as new datasets are generated, evidence may be found to support these interactions.

eY1H assays provide a powerful high-throughput PDI mapping method to test the binding of hundreds of TFs to defined DNA regions of interest. However, eY1H assays are not without caveats as they cannot detect TFs that exclusively interact with DNA as heterodimers, in cooperative complexes, or after post-translational modifications (Sewell and Fuxman Bass, 2018). In addition, eY1H-derived PDIs occur within the yeast nucleus, outside of the endogenous chromatin context which plays an important role in immune regulation (Smale et al., 2014). Improvements to the eY1H method, such as co-expressing two TFs or expressing phosphomimetic TF variants, and integration with chromatin accessibility and histone marks, will likely reduce the false negative rate and identify the appropriate cellular contexts in which the PDIs are relevant. The rate of false positives is conceptually more complicated to define as interactions can be specific to a cell-type or condition in which they have not been tested. Nonetheless, we found that interactions identified by eY1H assays and interactions that have been reported in the literature or identified by ChIP-seq validated at similar rates.

The eY1H dataset is enriched in NRs. NRs are ligand-activated TFs and thus they present an opportunity to modulate cytokine activity using drugs, especially in inflammatory diseases. Although antibodies have proven to be effective approaches in autoimmune diseases, approved antibodies blocking cytokine activity are available for only nine cytokines (DrugBank, (Wishart et al., 2018)). Additionally, a therapeutic strategy may require the induction of cytokine activity rather than inhibition, or the concomitant modulation of multiple cytokines. Thus, NRs may provide an alternative therapeutic approach to modulate cytokine expression. Here, we demonstrated the use of the eY1H and literature datasets as a framework to identify combinations of NR agonists that synergistically increase the production of IL10. Further exploration of the eY1H- and literature-derived datasets may identify other NRs that synergistically or concomitantly modulate the expression of multiple cytokine genes.

The eY1H data provides a resource of PDIs to integrate with other large-scale datasets and make functional predictions about TFs in cytokine regulation. Here, we present examples from integrating the eY1H data with scRNA-seq datasets to identify novel TFs and TF-cytokine regulatory axes in inflammatory diseases and lineage differentiation. Applying a similar approach, the eY1H data can be used to complement CRISPR screens to identify direct interactions between TFs and cytokine genes. Overall, the eY1H PDIs provides a powerful resource that can be mined in myriad additional ways to complement other datasets and delineate a more accurate understanding of cytokine gene regulatory networks.

## Acknowledgements

We thank members of the Fuxman Bass lab, Dr. Thomas Gilmore, and Dr. Trevor Siggers, for helpful discussions and critically reviewing the manuscript. We thank Joseph Bloom for assistance in retrieving the ENCODE and GTRD ChIP-seq datasets. We thank Callen Bragdon and Andrew Munoz for assistance in retrieving the NURSA Transcriptomine dataset. We thank Kyle Pedro for assistance in culturing PBMCs. This work was supported by US National Institutes of Health grants R00 GM114296 and R35 GM128625 to J.I.F.B., and R01 AI138960 and R01 AI122842 to A.H.J. L.M.A. was partially supported by 109263-59-RKRL, J.A.S. was partially supported by 5T32HL007501-34, and S.Y. was partially supported by Boston University Undergraduate Research Opportunities Program and New England Biolabs Summer Undergraduate Research Award.

## Author Contributions

C.S.S. and J.I.F.B. conceived the project, designed the experiments, and wrote the manuscript. C.S.S. and S.C.P. generated CytReg v2. J.I.F.B., C.S.S., and L.X., performed eY1H assays. C.S.S. and K.A.G. performed luciferase assays. C.S.S. and L.M.A. performed the PBMC experiments. S.L. and C.S.S. performed the iBMDM experiments. S.Y. and C.S.S. performed the THP-1 experiments. C.S.S., L.Z., J.I.F.B., S.C.P., and J.A.S., performed the data analysis. A.J.H. supervised L.M.A., and M.A. supervised S.L. All authors read and approved the manuscript.

## Figure Legends

**Supplementary Figure 1. Disease-associated PDI network**

eY1H disease-associated PDI network comprising 377 TF-cytokine-disease associations (edges) between 60 cytokines (yellow rectangle nodes) and 73 TFs (red circle nodes) associated with the same inflammatory diseases (DisGeNET, GWAS, or MGI). Edge color indicates the inflammatory disease that both the TF and cytokine are associated with.

**Supplementary Figure 2. Analyses of HIV scRNA-seq data**

(A) UMAP plots of cell clusters from HIV-infected subjects. A total of 1,102 dendritic cells (845 HIV+ and 257 healthy), 767 B cells (671 HIV+ and 96 healthy), and 2,679 T cells (2,508 HIV+ and 171 healthy), were obtained from the scRNA-seq data (Farhadian et al., 2018).

(B) Average expression and percentage of expression of cell calling markers in each cell population.

(C-E) Order of single cells along the HIV disease pseudotime trajectory from disease (dark blue) to healthy (light blue) for dendritic cells (C), B cells (D), and T cells (E), constructed using Monocle.

**Supplementary Figure 3. Analyses of IBD scRNA-seq data**

(A) UMAP plots of cell clusters from IBD-infected subjects. A total of 4,478 intestinal epithelial cells (1,436 inflammatory and 3,042 normal), 382 dendritic cells (274 inflammatory and 108 normal), 4,164 macrophages (2,316 inflammatory and 1,848 normal), 10,409 B cells (7,403 inflammatory and 3,006 normal), and 18,525 T cells (10,121 inflammatory and 8,404 normal), were obtained from the scRNA-seq data (Martin et al., 2019).

(B) Average expression and percentage of expression of cell calling markers in each cell population.

(C-G) Order of single cells along the IBD disease pseudotime trajectory from disease (dark blue) to healthy (light blue) for intestinal epithelial cells (C), dendritic cells (D), macrophages (E), B cells (F), and T cells (G), constructed using Monocle.

**Supplementary Figure 4. IBD pseudotime analyses**

TFs and cytokines in the IBD-associated PDI network that significantly change in expression along the pseudotime disease trajectory in each cell type. eY1H PDIs are connected by grey arches, and PDIs correlated/anti-correlated in expression on a single-cell basis in intestinal epithelial cells (brown edges), dendritic cells (orange edges), macrophages (pink edges), B cells (purple edges), or T cells (green edges), are indicated.

## Supplemental Information

**Table S1.** Literature-reported interactions between TFs and cytokine genes.

**Table S2.** TF associations with immune diseases and phenotypes.

**Table S3.** eY1H-derived interactions between TFs and cytokine genes.

**Table S4.** Functional evidence for eY1H-derived interactions involving nuclear receptors.

**Table S5.** TF and cytokine associations with inflammatory diseases.

**Table S6.** List TF-cytokine pairs in the HIV-associated PDI network correlated on a single-cell basis.

**Table S7.** List of TF-cytokine pairs in the IBD-association PDI network correlated on a single-cell basis.

**Table S8.** TF and cytokine associations with development of immune cell lineages obtained from DisGeNET, GWAS, and MGI.

**Table S9.** TF associations with immune cell differentiation or proliferative diseases obtained from the literature.

**Table S10. List of** TF and cytokine pairs in the immune cell lineage networks correlated on a single-cell basis.

**Table S11.** Supporting evidence for eY1H-derived PDIs.

**Table S12.** List of cytokine promoters tested by eY1H and luciferase assays.

**Table S13.** Sequences of shRNA guides used to generate iBMDM stable cell lines constitutively knocking down mouse TFs.

**Table S14.** Primers used to measure TF and cytokine expression by qPCR in human and mouse samples.

