## Supplementary figures and images for "Comprehensive mapping of the human cytokine gene regulatory network"

### Figure S1

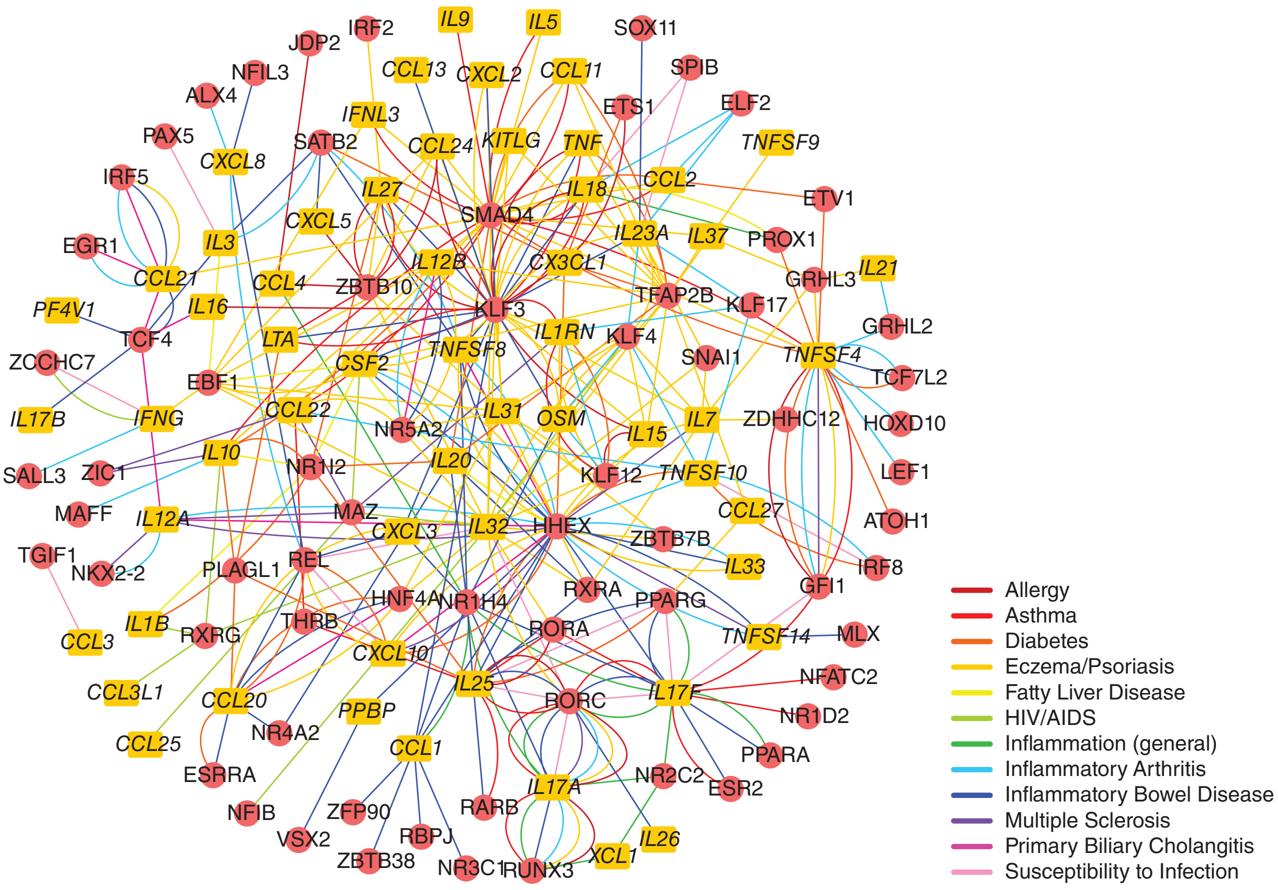

### Figure S2

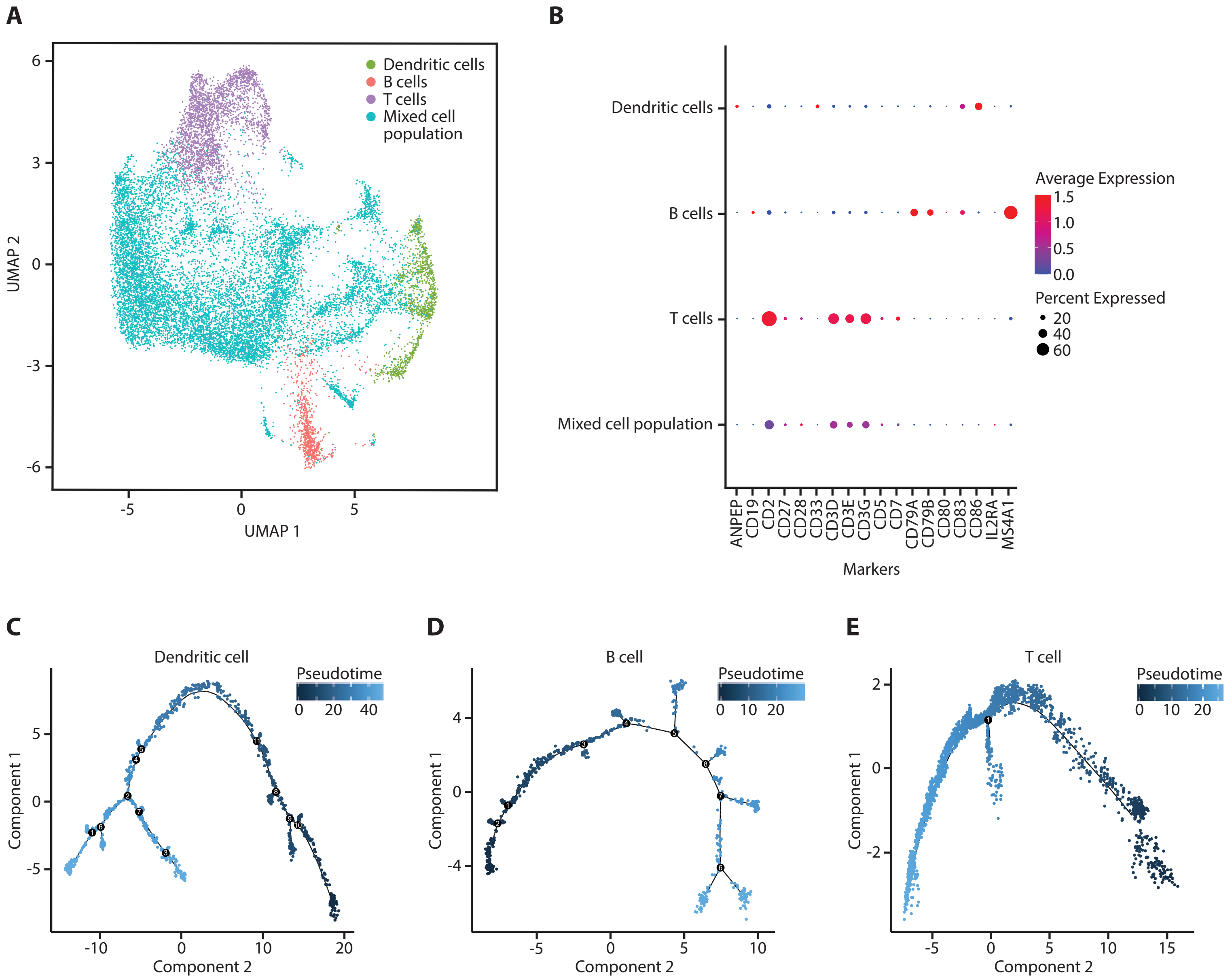

### Figure S3

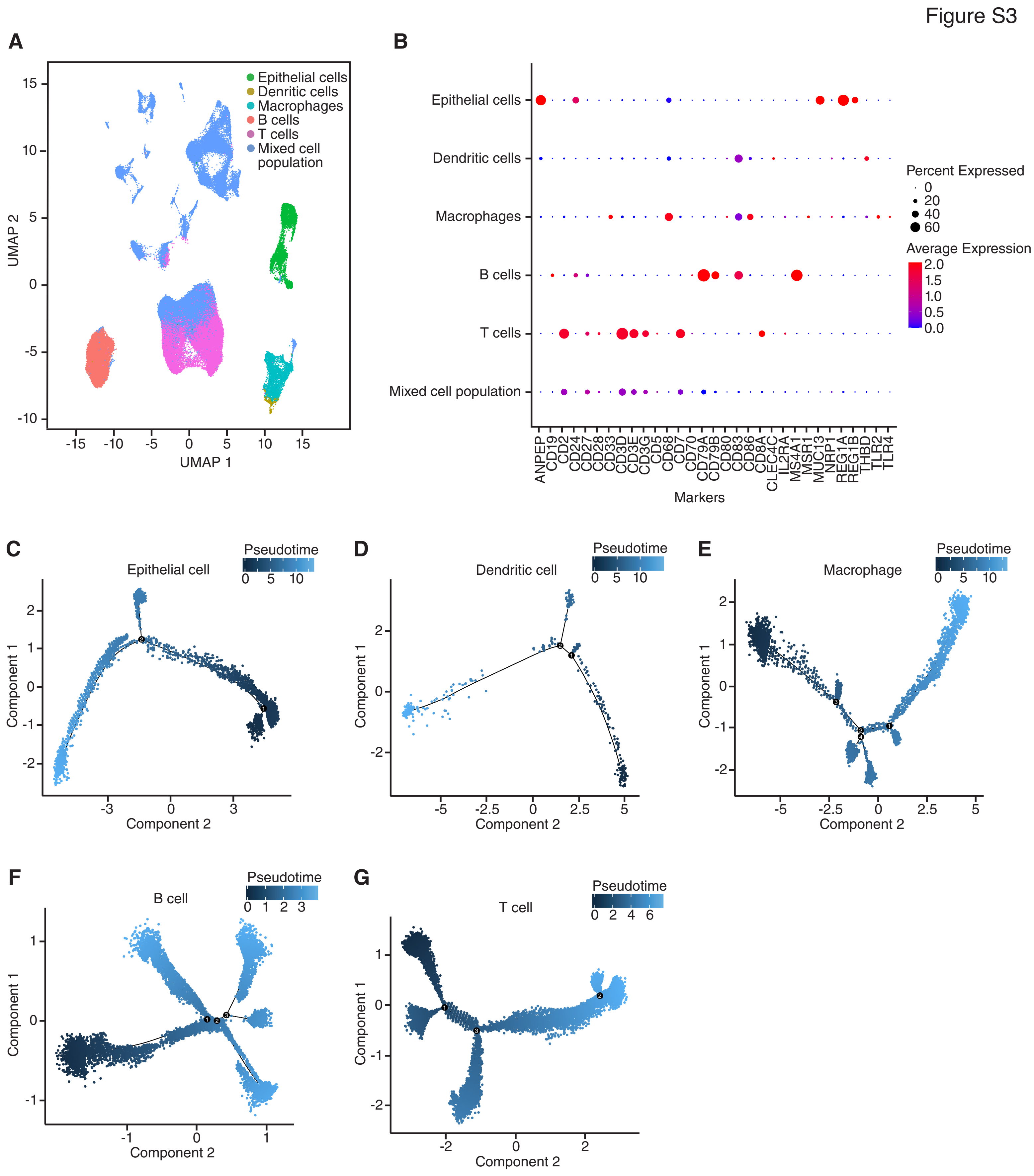

### Figure S4

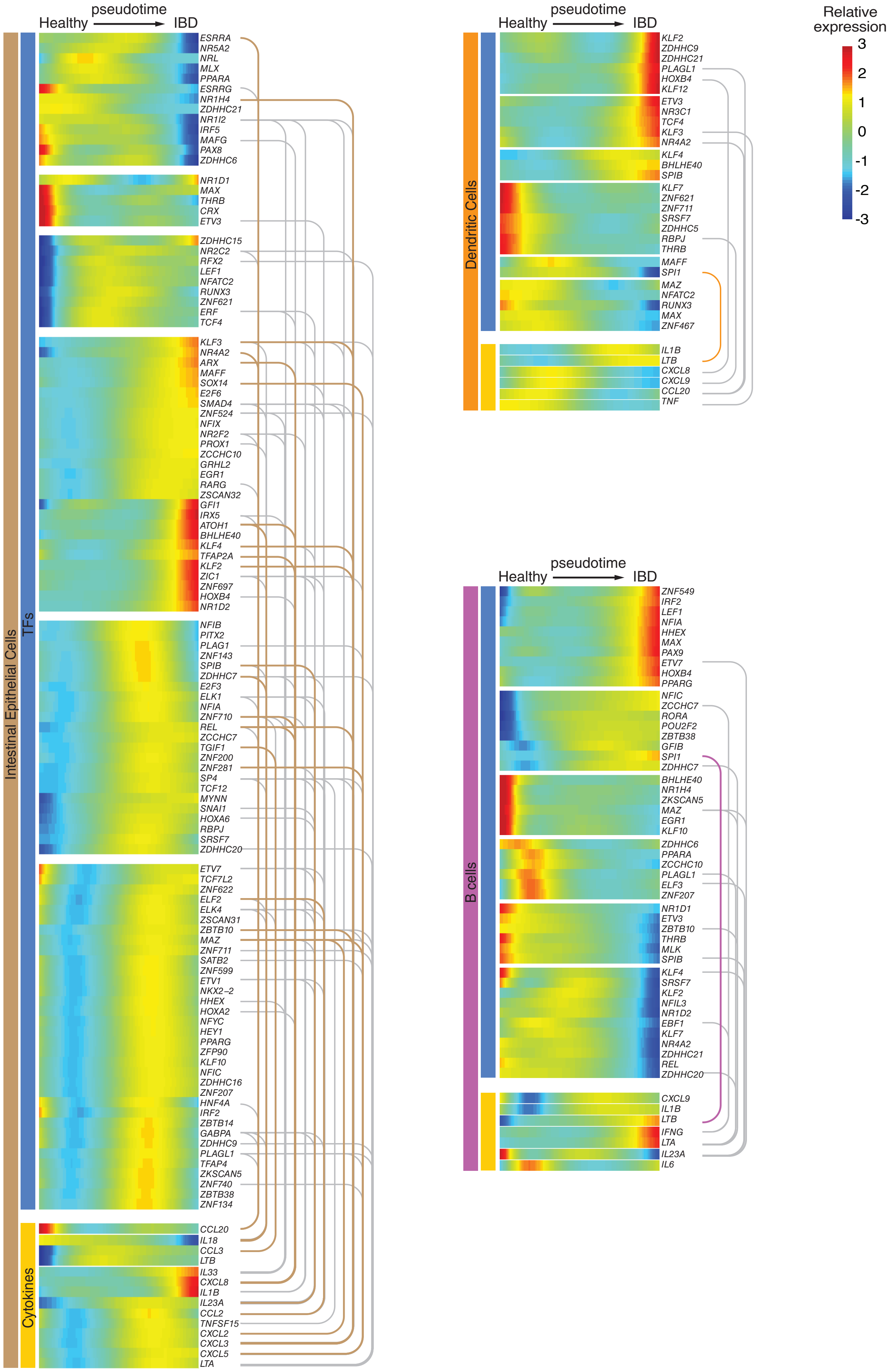

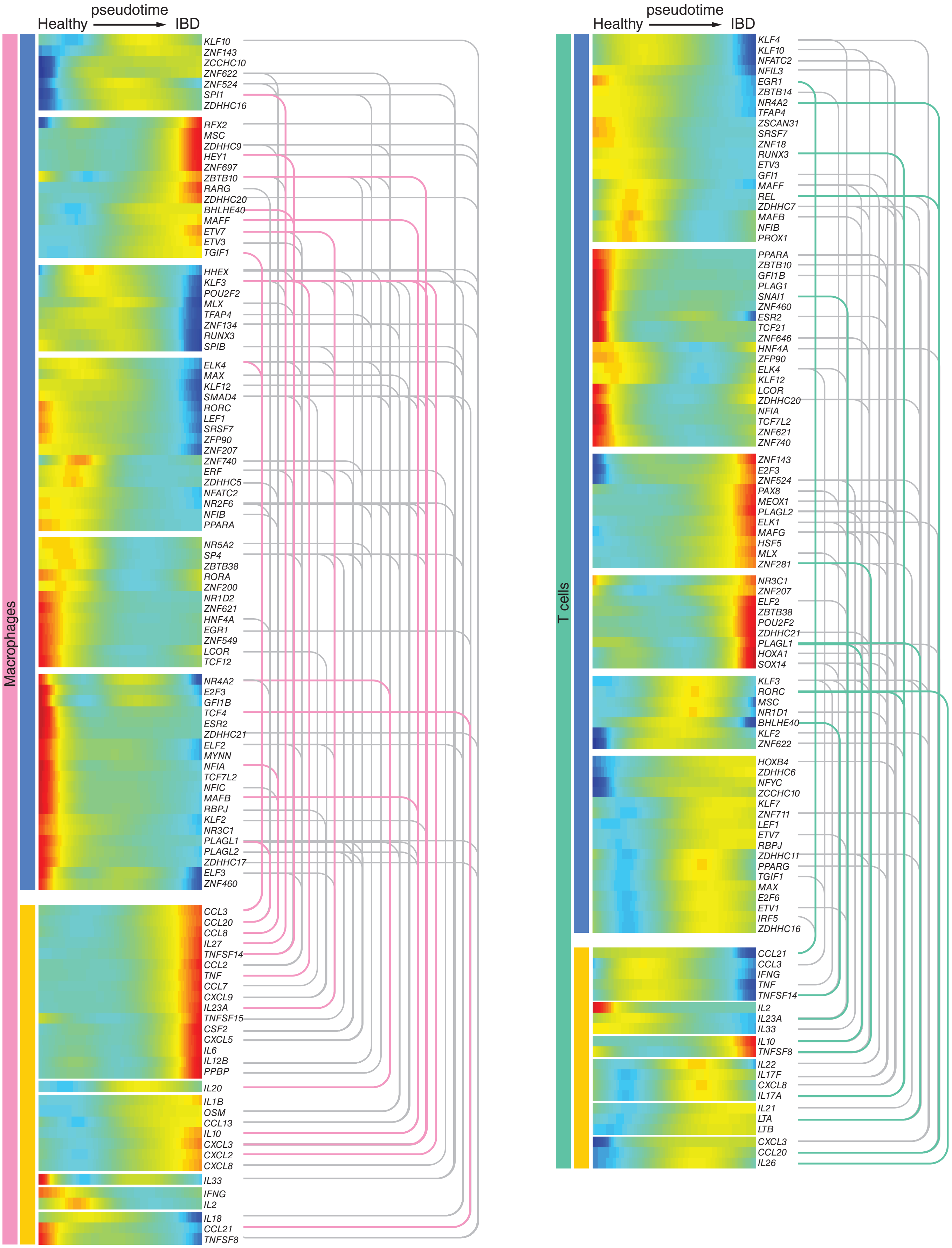
